# Tryptophan-Driven Metabolomic Shift in *Acidobacteriaceae* Reveals Phytohormones and Antifungal Metabolites

**DOI:** 10.1101/2025.10.02.680145

**Authors:** Celine M. Zumkeller, Christoph Hartwig, Walter Lanzalonga, Maria A. Patras, Yang Liu, Michael Marner, Sanja Mihajlovic, Marius S. Spohn, Till F. Schäberle

**Author notes:** Corresponding author: Till F. Schäberle.

## Abstract

Acidobacteriota is one of the most abundant phyla in soils and has recently attracted attention for its potential role in promoting phytosanitary benefits. The metabolomic capabilities of this phylum remain poorly characterised, with few experimentally confirmed metabolites described. To address these gaps, we combined metabolomic profiling with comparative genomic analysis to explore the functional potential of novel strains within the *Acidobacteriaceae* family. Genome mining across the phylum for plant-growth-promoting traits revealed the presence and taxon-specific enrichment of genes related to phytohormone production, as well as other genes associated with plant-beneficial microorganisms. When tryptophan was added to the cultivation medium, it triggered a strong metabolic response in the strains, resulting in measurable changes in the production levels of phytohormones such as indole-3-acetic acid and indole-3-pyruvate. Building on these findings, we examined the metabolic reprogramming caused by tryptophan supplementation, which suppressed the growth of phytopathogenic fungi, leading to the identification of malassezindoles and pityriacitrins as active agents. This was confirmed by isolating and elucidating the structure of pityriacitrin B and one of its methyl esters through NMR studies. Overall, these findings shed light on the previously unexplored metabolic potential of the Acidobacteriota phylum, emphasising its ecological importance for phytosanitary applications.

**Importance:** Despite their ubiquity and genomic diversity, the functional metabolism of members of the Acidobacteriota has largely remained uncharacterised. This study links genomic predictions to experimentally verified metabolomic outputs of Acidobacteriaceae, demonstrating tryptophan-responsive metabolic shifts translating to phytohormones and metabolites suppressing fungal growth. Our work underscores the emerging role of Acidobacteriota as important contributors to soil ecosystem functioning and plant–microbe interactions.

## Introduction

Ubiquitous across various ecosystems, including soil (1–3), marine sediments (4), wetlands (5–7), as well as extreme environments such as permafrost (8) and acidic mine drainages (9), Acidobacteriota represent one of the most widespread bacterial groups (10). The phylum is phylogenetically diverse (11), classified into 26 subdivisions (12) based on 16S rDNA sequencing. Despite their ubiquity, the Acidobacteriota have historically been underexplored due to the limited availability of cultured representatives, which has restricted experimental characterisation and genomic insights (13). Today, the availability of high-quality metagenome-assembled genomes, combined with the growing number of cultured representatives and isolate genomes, has significantly broadened our understanding of the metabolic capabilities and ecological roles of this phylum (13–15).

Initially described as slow-growing oligotrophs (16) With limited metabolic dynamics, adapted to nutrient-depleted environments, modern insights now reveal their versatility and critical involvement in biogeochemical processes, including sulphur (17) and nitrogen (18) cycling as well as H_2_ consumption (19). Furthermore, they can degrade plant-derived polysaccharides using their rich repertoire of carbohydrate-active enzymes, such as cellulases, chitinases, and xylanases (20). Finally, early studies dating back to 2006 proposed a substantial potential for secondary metabolite production by biosynthetic gene clusters, identifying novel Polyketide synthase gene clusters of acidobacterial origin from soil (21, 22). Subsequent genomic (23) and metagenomic studies confirmed this finding, identifying biosynthetic-gene-cluster (BGC)-enriched lineages such as the *Acanthopleuribacteraceae* (24) and the genus *Candidatus Angelobacter* (22). Yet, translating bioinformatic findings into experimental validation remains difficult and still requires the availability of culturable representatives. Notably, Leopold-Messer *et al.* (24)successfully retrieved marine sponge symbiont strains from the *Acanthopleuribacteraceae* family, marking the first description of non-ribosomal peptide (NRP)-polyketide (PK)-derived secondary metabolites from this phylum (24).

To overcome these gaps, various cultivation approaches have been developed to selectively favour acidobacterial growth and retrieve single-isolate strains from complex environmental samples (3, 25–27). These approaches have significantly increased the number of isolated strains; however, they still only represent a fraction of the predicted phylum diversity. Most of the strain isolates are retrieved from soil (23), where Acidobacteriota have consistently shown high abundances, ranging from 20% to 40% of the detected 16S rRNA(28, 29) sequences. Cultivation strategies often utilise plant polymers for Acidobacteriota enrichment (25, 26, 30), providing insight into their predicted function in enhancing soil health by degrading undigestible plant polymers and improving nutrient content in the soil. Consequently, several studies investigate the potential plant-growth-promoting (PGP) effects of the phylum. In 2016, the first evidence of a direct plant-interaction was described by Kielak *et al*. (31). They found that some acidobacterial strains produced IAA and also demonstrated a PGP effect on the model plant *Arabidopsis thaliana* (31). Moreover, Acidobacteriota are recognised for synthesising hopanoids (32), bacterial analogues of sterols, known to enhance stress tolerance in soil bacteria as an integral component of the bacterial membrane. Their unique exopolysaccharide production (33, 34) further suggests a potential ability to endure dry conditions and possible roles in sustaining plant health, e.g., protecting against desiccation. Complementary metagenomic and meta-transcriptomic studies have revealed a significant enrichment of genes related to PGP. Gonçalves *et al.* (2024) profiled 758 soil-associated metagenome-assembled genomes (MAGs) and identified plant-growth-promoting-traits (PGPTs) involved in phytohormone production, nitrogen fixation, phosphorus solubilisation, siderophore production, and exopolysaccharide biosynthesis (20). These traits were explicitly enriched in certain families of the Acidobacteriota, namely *Acidobacteriaceae*, *Pyrinomonadaceae*, and *Koribacteriaceae*. Meta-transcriptomic evidence shows that these traits are actively transcribed in their respective environmental context (20). Furthermore, transcriptomic research has demonstrated the metabolic dynamics and responsiveness to external stimuli of cultured Acidobacteriota strains. For instance, *Acidobacterium capsulatum* responds to oxygen deprivation by switching to a respiro-fermentative state, resulting in the production of acetate and ethanol —a strategy that allows for survival in habitats with fluctuating O_2_ levels, such as soil (35). The oligotrophic lifestyle typical of the Acidobacteriota has been investigated in a *Granulicella* species, which responds to high-carbon availability by upregulating stress-related genes, indicating sensitivity to nutritional changes (36). While many ecological functions are inferred from meta(transcriptomics) data, experimentally generated metabolomic data remain limited, primarily focusing on the characterisation of membrane lipids and their production in response to different cultivation parameters such as temperature, pH, and O2 (37, 38).

To address these gaps, we integrated metabolomic profiling with comparative genomic analysis to investigate the functional potential of novel strains of the *Acidobacteriaceae* family. Phylum-wide genome mining for PGPTs revealed the presence and taxon-specific enrichment of genes associated with phytohormone production, as well as additional genes related to PGP effects. The addition of tryptophan (Trp) to the cultivation medium elicited a strong metabolic response in the strains, resulting in quantitatively determined shifts in the production levels of the phytohormones indole-3-acetic acid (IAA) and indole-3-pyruvate (iP). Prompted by these findings, we evaluated the metabolic reprogramming induced by Trp supplementation, revealing suppressive effects on the growth of phytopathogenic fungi, and subsequently identified pityriacitrins and malassezindoles as the causative agents. Collectively, these findings highlight the previously underexplored metabolic space of the Acidobacteriota phylum, underscoring its ecological significance for phytosanitary purposes.

## Results and Discussion

### Result 1 - Diverse Cultivation Strategies Reveal the Production of Phytohormones

Extensive metagenomic data from globally distributed bioresources indicate an underexplored biosynthetic and ecological potential of the Acidobacteriota phylum. During our ongoing efforts to isolate and bioprospect bacterial diversity for its potential in the production of natural products and related environmental functions, we retrieved three Acidobacteriota strains from termite nest material (25) and one strain from a German soil sample (39). To evaluate their biosynthetic potential, we isolated their total DNA together with that of the type strain *A. borealis* DSM 23886 and performed whole-genome sequencing. The generated genomic data underwent quality check using CheckM2 (40). By applying recognised quality criteria (completion >90% and contamination <5% (41)), one assembly (FHG110206) was disqualified for further analysis, while the other three assemblies were used for genome-based taxonomic affiliation by the genome taxonomy database (GTDB) (42) (Supplementary Table 1). This positioned the termite nest-associated strains as members of the genera *Acidobacterium* (FHG110202) and *Terracidiphilus* (FHG110214), while the soil-associated strain FHG110511 was identified to be an *Edaphobacter*. All three strains belong to the family of *Acidobacteriaceae* (collectively referred to as FHG *Acidobacteriaceae* in this manuscript). Thus, we selected further available strains of this family (*Edaphobacter aggregans* DSM 19364*, Acidobacterium sp.* S8*, Silvibacterium bohemicum* S15*, Acidicapsa borealis* DSM 23886*, Granulicella rosea* DSM 18704, and *Bryocella elongata* DSM 22489)(Table S1) and performed an extensive OSMAC (one-strain-many-compounds) approach (43), conceptualised to increase the likelihood of covering a favourable condition for the strains to proliferate and promote gene expression. To this end, we modified the carbon sources, vessel size, and shaking speed, and applied solidification and surface texture methods using quartz or ceramic beads, potentially enabling the surface attachment of the cells. Cultivations were performed on a 100 ml scale using 300 ml Erlenmeyer flasks or at a 4 ml scale using 24 well Duetz system plates (Table S2). Samples were taken on days 5, 7, 10, 12, and 14 of the respective cultivations. The generated samples were dried by lyophilisation and extracted with methanol, resulting in a total of 892 organic extracts. An aliquot of each extract was reconstituted in dimethyl sulfoxide (DMSO) and used to determine the antimicrobial bioactivity by screening against the bacterial indicator strains *Escherichia coli* ATCC 35218, *Bacillus subtilis* DSM 10, and *Mycobacterium smegmatis* ATCC 607, as well as the fungus *Candida albicans* FH2173. This, however, did not reveal any reproducible antimicrobial effect enabling traceability of a causative agent (data not shown). Furthermore, we conducted a bioactivity-independent and metabolomics-driven bioprospecting campaign. Initially, we focused on the metabolic profiling of strains FHG110202 and FHG110206 and assessed the metabolic composition of their generated extracts (n = 244, including media controls) analytically by performing ultra-high-performance liquid chromatography coupled to high-resolution mass spectrometry (UHPLC-QTOF-HR-MS). Molecular features (*m/z*, retention time) were detected and aligned into buckets. Combining timepoints for each strain-media combination and presence-absence analysis led to 72 conditions (including media controls) with 20285 features present and distributed into 2747 buckets (Figure S1).To assess the effect of the medium on the metabolome of the strains, we filtered medium-derived features (n=1795). Pairwise Jaccard distances between samples were ordinated by principal coordinates analysis (PCoA) for strains FHG110206 and FHG110202 (Figure 1A). Metabolomic profiles clustered mainly by medium (colour code) and vessel size (top, left cluster) rather than by strain. We also observe a distinct cluster for medium containing quartz beads (bottom, right cluster). A PERMANOVA analysis confirmed this observation, suggesting a strong OSMAC response. The medium identity accounts for 87% of the variance among strain extracts (F=7.09, p=0.001), while the strain identity within the same medium explains a much smaller share (1.7%, F=0.77, p=0.001). Within the small-scale set, the bead condition explained 29.2% of the differences (F = 7.64, p = 0.001), consistent with the observed quartz-bead cluster. Thus, metabolomes are primarily influenced by the medium and vessel size, with only a small strain-level difference. We also plotted unique features of strain-produced metabolites for individual media conditions (Figure 1B), and found that in both strains (FHG110202:FHG110206), media F (87:125), U (65:64), W (86:86), and X (197:116) are enriched in unique features compared to the average unique feature number across all media (22.2:20.7). This indicates that the large-scale cultivation size in 300 ml Erlenmeyer flasks yields a more distinct metabolome compare to the small-scale cultivation in 24-well plates.

**Figure 1.**
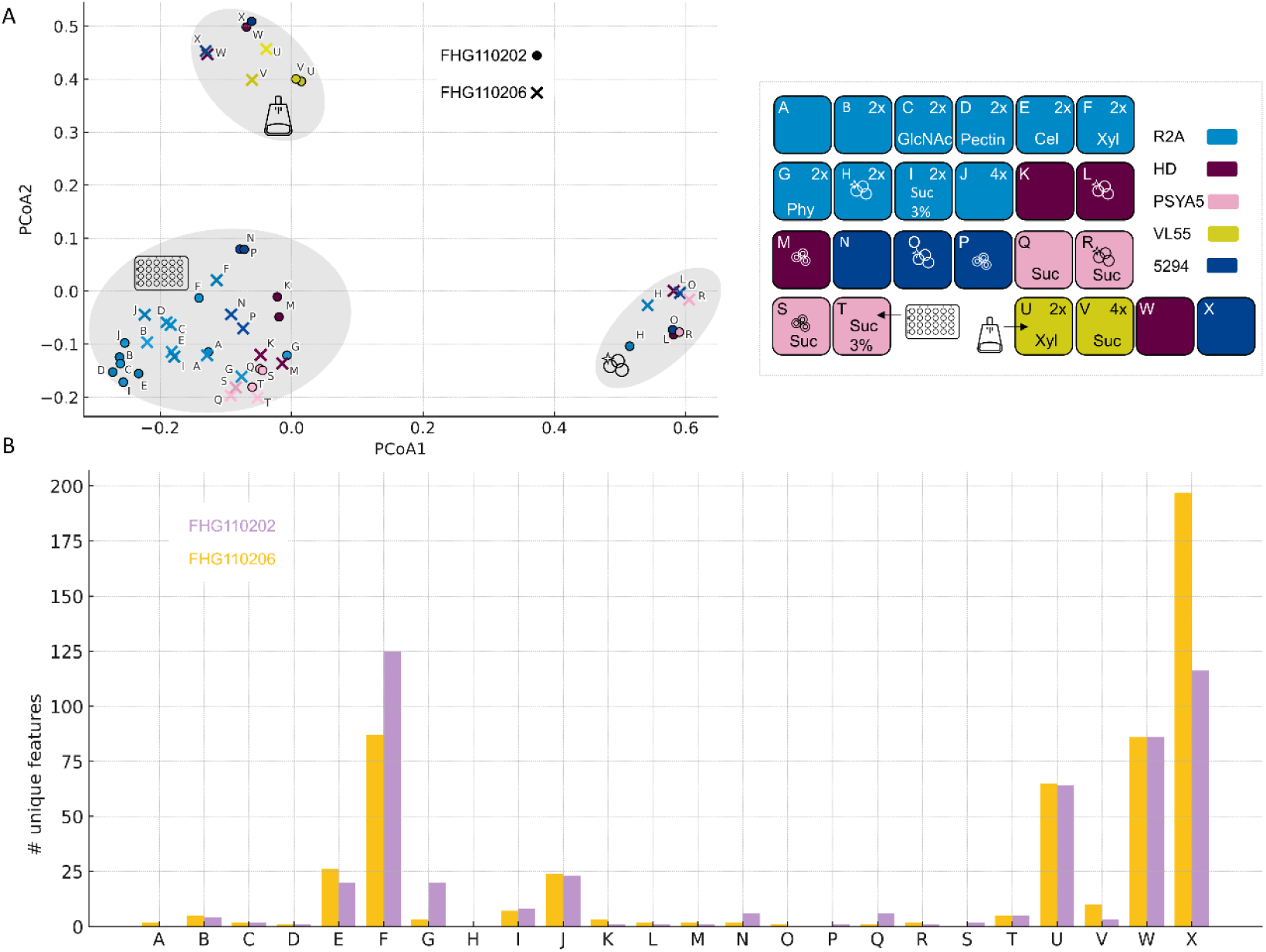
OSMAC metabolome variation by medium, strain, and cultivation scale. The legend details the media compositions (A-X), based on base-media colour, concentration, additives, and bead status. Samples labelled A - T were cultivated in 24 - well plates (small-scale), while U-X were grown in 300 ml Erlenmeyer flasks. I f not stated otherwise, additives were supplied in concentrations of GlcNAc, N - acetyl-glucosamine (0.05 %); Cel, cellobiose (0.5 %); Xyl, xylan (0.5 %); Phy, phytagel (0.8 %); Suc, sucrose (0.5 %). **(A) PCo A of control-filtered LC/ MS presence/ absence fingerprints using Jaccard distance**. Samples are labelled by the medium letter (A - X) and coloured by base-medium (see legend). Samples from FHG110202 are marked with circles, and FHG110206 with crosses. Light grey ellipses highlight visually distinct clusters, and icons indicate their common features: large - scale flask cultivation (flask), small - scale 24 - well cultivation (24 - well plate), and quartz beads containing samples (circles). **(B) Number of unique features per medium (A - X) and strain as clustered bars**. Strains are indicated by colour: FHG110202 (light purple), FHG110206 (yellow). Unique features are defined as not being present in any medium control and only present in the respective medium, but not in any other medium.

The detected metabolite features were matched against our in-house reference compound MS database, which contained approximately 1300 structurally characterised microbial metabolites at the time of data processing. By comparison of mass to charge ratios and retention time, we were able to annotate the phytohormone IAA in the extracts of our FHG *Acidobacteriaceae* strains with a confidence level 1 (Identified metabolite: as proposed by the Chemical Analysis Working Group of the Metabolomics Standards Initiative (44)). The production of the phytohormone IAA has been reported for members of the *Acidobacteriaceae* (31). To systematically describe the cultivation conditions that favour IAA production, we visualised the peak area of IAA in FHG110202 as an example across all tested media and setups (Figure S2). While IAA was not produced under conditions leading to the isolation of the strains (VL-55 base media), it became detectable in more complex media. Among these, R2A-based cultivation, with agitation, consistently showed the best IAA production. To further analyse the phytohormone production of FHG110202, we included additional standards: Kinetin (CAS 525-79-1), Trans-zeatin (CAS 1637-39-4), Indole-3 Butyric Acid (CAS 133-32-4), Brassinolide (CAS 72962-43-7), jasmonic acid methyl ester (CAS 39924-52-2), jasmonic acid (CAS 77026-92-7), abscisic acid (CAS 21293-29-8), indole-3-propionic acid (CAS 830-96-6), 1,2-Diphenylurea (CAS 102-07-8) and *N6*-(*Δ2*-Isopentenyl)adenine (iP)(CAS 2365-40-4) to our *in-house* reference database and explicitly checked for their presence in the generated crude extracts. This revealed the presence of the cytokinin iP at a confidence level of 1. The detection of cytokinin iP as a biosynthetic product of members of the Acidobacteriota phylum is a new finding.

### Result 2 – Biosynthetic Potential of the Acidobacteriota phylum

To complement the metabolomic workflow, we evaluated the genomic potential of our FHG *Acidobacteriaceae* and comparatively aligned it to the broader Acidobacteriota landscape. Therefore, we initially searched for publicly available genomes matching the criteria Query: “Acidobacteriota” NOT “anomalous” in the NCBI database. This yielded 1428 genomes that underwent quality control using CheckM2 (40) (Figure S3), applying the above-defined quality criteria. This resulted in a final dataset comprising 618 genome assemblies. Overall, most of the analysed genomes were retrieved from metagenomes (n = 533; 86%) (Supplemental Table 1). A detailed bioinformatic analysis of this dataset is reported in the Extended Methods and Results section of the supplementary information. The taxonomic affiliation was established by GTDB (42) and the generated multiple-sequence alignment was used to build a phylogenetic tree. The bioinformatically predicted features of each strain were assigned and visualised within the tree using iTOL (45). The genomes in the dataset span 12 classes (Figure 2), dominated by the *Acidobacteriae* (n = 242), *Blastocatellia* (n = 130) and *Vicinamibacteria* (n = 98), which also represent the majority of today’s strains brought to culture (76/86-88%). To evaluate the biosynthetic potential of the Acidobacteriota phylum, we predicted BGCs using the standalone version of antiSMASH (46). In total, 3865 BGCs were detected in the genome dataset. The most abundant BGC class in the dataset is ribosomally produced and post-translationally modified peptides (RiPPs) (n = 1152, 30%), followed by non-ribosomal peptide synthetases (NRPSs) (n = 978, 26%) and terpenes (n = 777, 13%). The total average cluster number per class in our acidobacterial dataset is 6.2 ± 4.4 BGCs. The *Acidobacteriae* class exhibits the highest overall number of BGCs, with an average of 7.4 ± 3.7 BGCs per genome, followed by UBA6911 and Mor1, with 6.3 ± 2.5 and 5.8 ± 2.1 BGCs, respectively. Dedicated analysis of the FHG *Acidobacteriaceae* revealed an average of 9 ± 3.46 BGCs, slightly higher than for the overall phylum and the *Acidobacteriae* class value, but comparable to the overall family value (7.9 ± 3.0). Again, the most represented group of BGCs is the RIPPs (sum: 10), followed by the terpenes (sum: 7) and the NRPS (sum: 4) BGCs. Notably, the *Holophagae* class, despite having an average BGC number of 5.1, exhibits a high standard deviation of 10.5, indicating a high potential for some of its representatives. This value can be accounted for by the *Acanthopleuribacteraceae* (Z-score: 3.5), showing high BGC counts of >40, seven times higher than the average Acidobacteriota genome. Intriguingly, the first acidobacterial NPs recently isolated also come from this family (24). While these talented strains have been isolated from marine animals, Acidobacteriota are known to be habitat generalists with a wide distribution, primarily found in soil ecosystems. They are characterised by a variety of extracellular products, such as biosynthetic gene clusters (BGCs), which are more extensive in soil-preferring lineages compared to their non-soil-preferring relatives, including the *Holophagae*. Although analysing biosynthetic potential at the class level is useful, our analysis demonstrates the advantages of examining family and genus levels. This approach helps identify the most promising taxonomic hotspots for discovering new natural products (NP).

**Figure 2:**
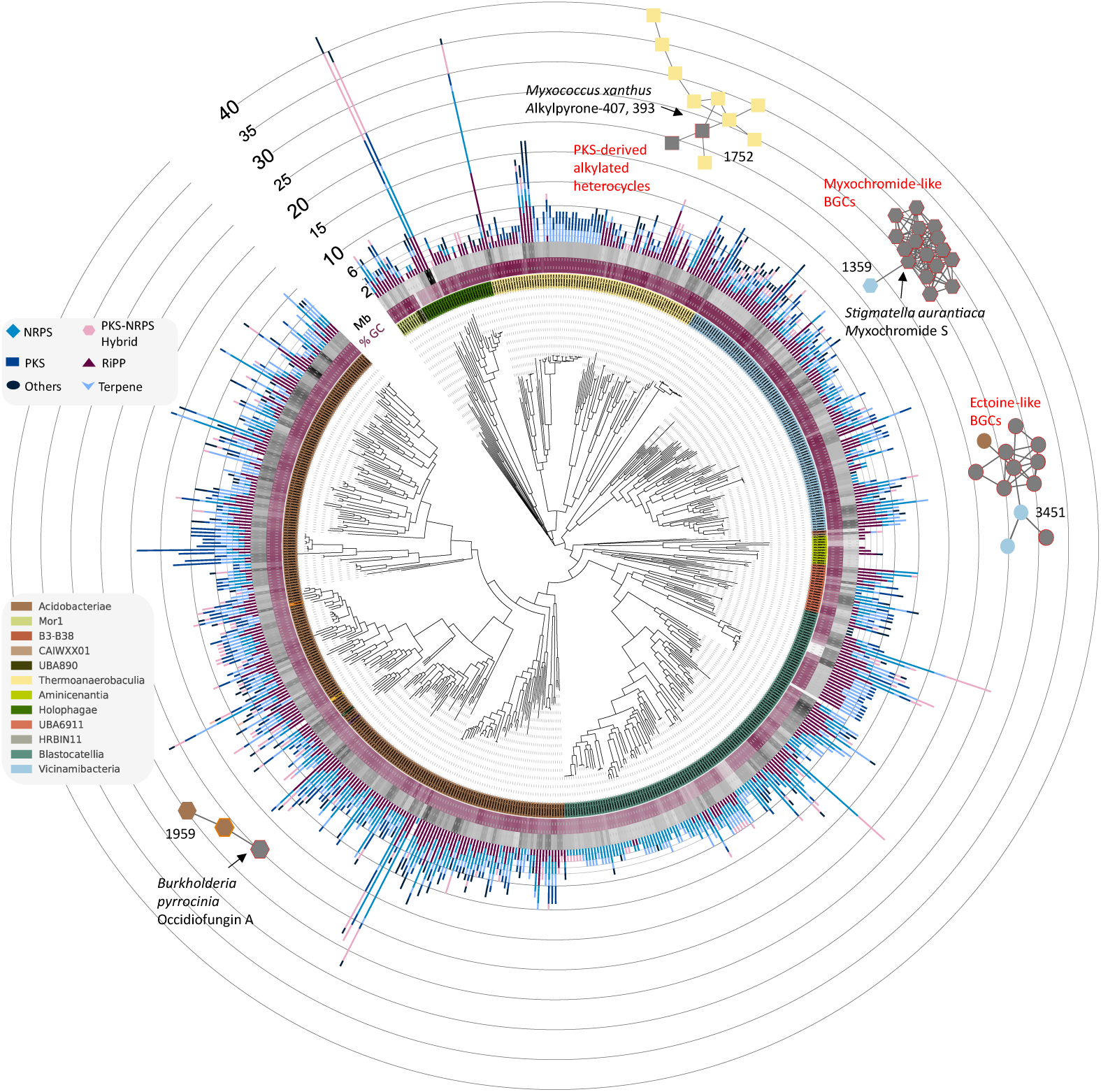
BGC composition of the Acidobacteriota Phylum and Bi G-SCAPE Network of acidobacterial BGCs clustering with MiBIG reference clusters. Multiple Sequence Alignments generated by GTDB were used to create a phylogenetic tree of 618 representatives of the Acidobacteriota. Twelve acidobacterial classes were identified by GTDB and are indicated by different colours (ring 1). FHG strains are colour-coded as follows: FHG110214 (green), *A. borealis* DSM 23886 (purple), Fh G110202 (yellow), and Fh G110511 (orange). Ring two indicates the genome size of the individual representatives, ranging from black (larger genomes) to grey (smaller genomes). Bar charts indicate the total number of antiSMASH - detected BGCs, differentiated into different BGC types.

To complement the evaluation of the Acidobacteriota phylum, we performed sequence similarity calculations on the total of 3865 BGCs in our dataset using BiG-SCAPE (47), including the MiBIG reference BGCs, and applied a cutoff of 0.5. 1651 BGCs remained as singletons, contrasted by the largest Gene Cluster Family (GCF) that consists of 25 members, all from the *Acidobacteriae* class. Twenty-five GCFs are specific to one respective taxonomic class (Figure 1B, Figure S8); most of these GCFs contain more than 10 representatives (16/25), indicating clear taxonomic conservation. This taxonomic conservation is further illustrated by comparing the GCF composition of the different strains using cosine distance and visualising this similarity as uniform manifold approximation and projection (UMAP) (Figure S9). Here, strains belonging to the same taxonomic classes cluster within the visualisation, indicating overall conservation between phylogenetic groups. However, sequence similarity calculations revealed that the FHG *Acidobacteriaceae* do not share any of their clusters (Figure S8). Further, alignments with MiBIG Reference BGCs and confirmation by CORASON alignment (47) revealed a co-clustering of 7 acidobacterial BGCs in four GCFs (Figure 2), showing compositional BGC similarity to, e.g. the myxochromide and the occidiofungin A-reference BGCs (SI-extended methods and results). Intriguingly, the latter is a match to our strain FHG110511, harbouring a PKS/NRPS hybrid BGC that clusters in GCF1959 with the occidiofungin A reference BGC (MiBIG BGC0001711) at a pairwise identity of 70.4% (Figure S11). Occidiofungin A is a *Burkholderia contaminans-*produced NP that exhibits broad antifungal activity and is in early clinical development (gel OCF001, phase 1) for use against multidrug-resistant vaginal yeast (*Candida* spp.) infections (48, 49). The comparative evaluation of the NRPS and NRPS/PKS hybrid core gene architecture revealed a highly conserved module arrangement with certain discrepancies in predicted A-domain specificity. Testing of the strain’s organic extracts against *C. albicans,* however, showed no antifungal activity.

The occidiofungin-like BGC represents a singleton hit in respect to multimodular NRPS, PKS or hybrid systems within the genomes of the FHG *Acidobacteriaceae*, indicating a generally underdeveloped talent of our strains to produce structurally complex NPs and a possible reason for the lack of antimicrobial activity observations during our initial bioprospecting campaign.

### Result 3 – Tryptophan Supplementation Increases IAA but Decreases iP Production in Acidobacteriaceae without Enhancing Barley Seedling Biomass

During the metabolomic-based inspection of the OSMAC experiment, we identified the biosynthetic potential of our FHG *Acidobacteriaceae* to produce the phytohormones IAA and iP. To further profile the IAA and iP production of our FHG *Acidobacteriaceae* and the reference strain *A. borealis* DSM 23886, we cultivated those strains in media with and without supplementation of tryptophan (Trp), commonly known to increase IAA production in microorganisms (50). We performed a kinetics study to analyse the influence of 0.2% Trp on the relative IAA and iP production over 14 days following the methanolic extraction of the cultures. Overall, the amounts of IAA and iP increased over time, reaching their highest levels on days 10 and 12 of growth (Figure 3 and S12). Indeed, the supplementation of Trp enhanced IAA production significantly in all strains. In contrast, supplementation of Trp consistently suppressed iP production in the analysed strains except for FhG110202 and *A. borealis* DSM 23886, where this effect is less pronounced (Figure 3A). This initial profiling revealed that strain FHG110214 was the most productive iP producer, while exhibiting comparable productivity in IAA to *A. borealis* DSM 23886, FHG110202, and FHG110206. Based on this production profile, we selected FHG110214 and *A. borealis* DSM 23886 to perform an absolute quantification of phytohormones. When supplemented with Trp, *A. borealis* DSM 23886 produced IAA to a final concentration of 0.326 ± 0.063 μg/ml whereas FHG110214 produced IAA at a concentration of 0.104 ± 0.021 μg/ml. Instead, without supplementation of Trp, the IAA titres dropped to 0.124 ± 0.029 μg/ml (3-fold) and 0.015 ± 0.003 μg/ml (7-fold) in *A. borealis* DSM 23886 and FHG110214, respectively. This production profile is reversed for iP, where Trp supplementation resulted in decreased production for both strains FHG110214 (With Trp: 0.0058 ±0.0012 μg/ml; without Trp: 0.0143 ± 0.0024 μg/ml) and *A. borealis* DSM 23886 (With Trp: 0.005 ±0.0004 μg/ml; Without Trp: 0.0108 ± 0.0008 μg/ml). These production titres are in a comparable range to quantities previously reported for plant-growth-promoting bacteria (PGPB) (51, 52) and, e.g. 10-fold lower compared to those reported for *Pseudomonas*, *Streptomyces*, and *Enterobacter* species, which have been shown to produce 10–150 µg/ml under optimised conditions. The production of the cytokinin iP has not been described so far for the Acidobacteriota phylum. We quantified its production to 0.003 µg/ml in the most productive strain of our study. This is in alignment with the literature, which reports, for example, 17.5 pmol/ml (translating to 0.00356 µg/ml) in *Achromobacter xylosoxidans* (53) and lower concentrations such as 0.0003 µg/ml (1.5 nM) in *Paenibacillus polymyxa* (54).

**Figure 3:**
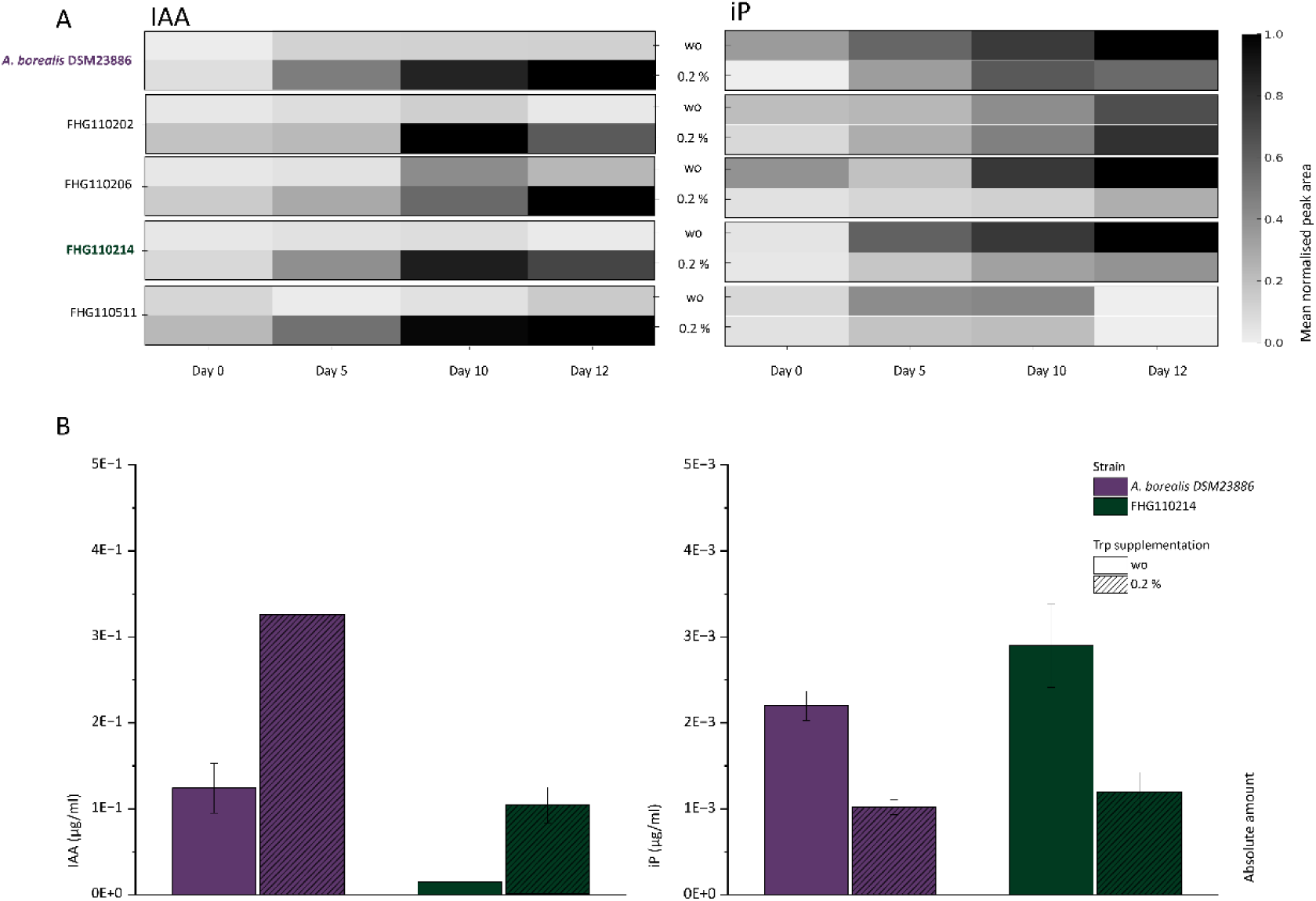
Relative quantification of IAA and iP in FHG *Acidobacteriaceae* and *A. borealis* DSM 23886 as normalised peak area (A) and absolute quantification of these phytohormones in the top producers FHG 110214 (iP) and *A. borealis* DSM 23886 (IAA)(B). **A** Raw peak areas were normalised against the largest peak area within one biological kinetic study for IAA and iP separately. Mean values were calculated and plotted as a heatmap to indicate temporal and dynamic induction patterns between strains and supplementation status. Trp supplementation results in higher IAA production but generally has a negative impact on iP production. This effect i s less pronounced in FHG110202 and *A. borealis* DSM 23886. **B** Absolute concentrations were derived from a calibration curve of IAA and iP, correlating peak area with given concentrations. Cultivations were repeated in triplicate, and the average of the calculated concentrations is shown for *A. borealis* DSM 23886 (green) and FHG110214 (purple).

Previously, it has been shown that culture broths of IAA-producing *Acidobacteriaceae* promoted root and shoot growth in *Arabidopsis thaliana* (20, 31) and growth and chlorophyll content in duckweed (55). Furthermore, cytokines have also been shown to have a beneficial impact on plant growth and defence response against pathogens and pests (56). To evaluate the plant growth-promoting potential of our extracts, we applied them in concentrations matching the culture concentration on barley seedlings. Additionally, we included a media extract control, as well as IAA and iP at defined concentrations, for testing. Briefly, extracts were spotted into glass vials containing 5 ml SH-medium, and the solvent was evaporated before one barley seed was added per vial. After an initial phase of darkness for germination, a day/night cycle incubation was performed for 6 days (Figure S13). Afterwards, the plants were extracted from the medium, cleaned, and separated into root and shoot before being lyophilised for dry biomass determination (Figure 4A and B). All samples showed a consistently higher biomass if the sample contained Trp. The pure compound controls showed growth comparable to that of the medium with IAA at a concentration of 0.442 μM, resulting in a biomass of 11 mg (similar to the medium control containing 0.2% Trp). Compared to the controls, extracts from FHG110214 and *A. borealis* DSM 23886 decreased plant biomass in the shoots (Figure 4A) and roots (Figure 4B) with and without added Trp. However, the addition of Trp to the cultivations decreased the adverse growth effect of the extract itself. Overall, the crude extracts of *A. borealis* and FHG110214 did not significantly promote plant growth of barley seeds in the experimental setup tested. It has been demonstrated that IAA can promote plant growth when bacterial colonies are in direct contact with roots (31). Our experimental setup, in contrast, was executed using cell-free organic extracts to provide a controlled sample. This, however, may not replicate localised, high-concentration phytohormone effects occurring in natural rhizosphere environments. Furthermore, applying cells or cell-free supernatants may provide further traits, such as enzymes with a beneficial effect on plant growth, not covered by organic extracts. In future studies, live bacterial cultures or co-cultivation systems with plants would provide more ecologically relevant data, albeit with increased challenges associated with cultivation.

**Figure 4:**
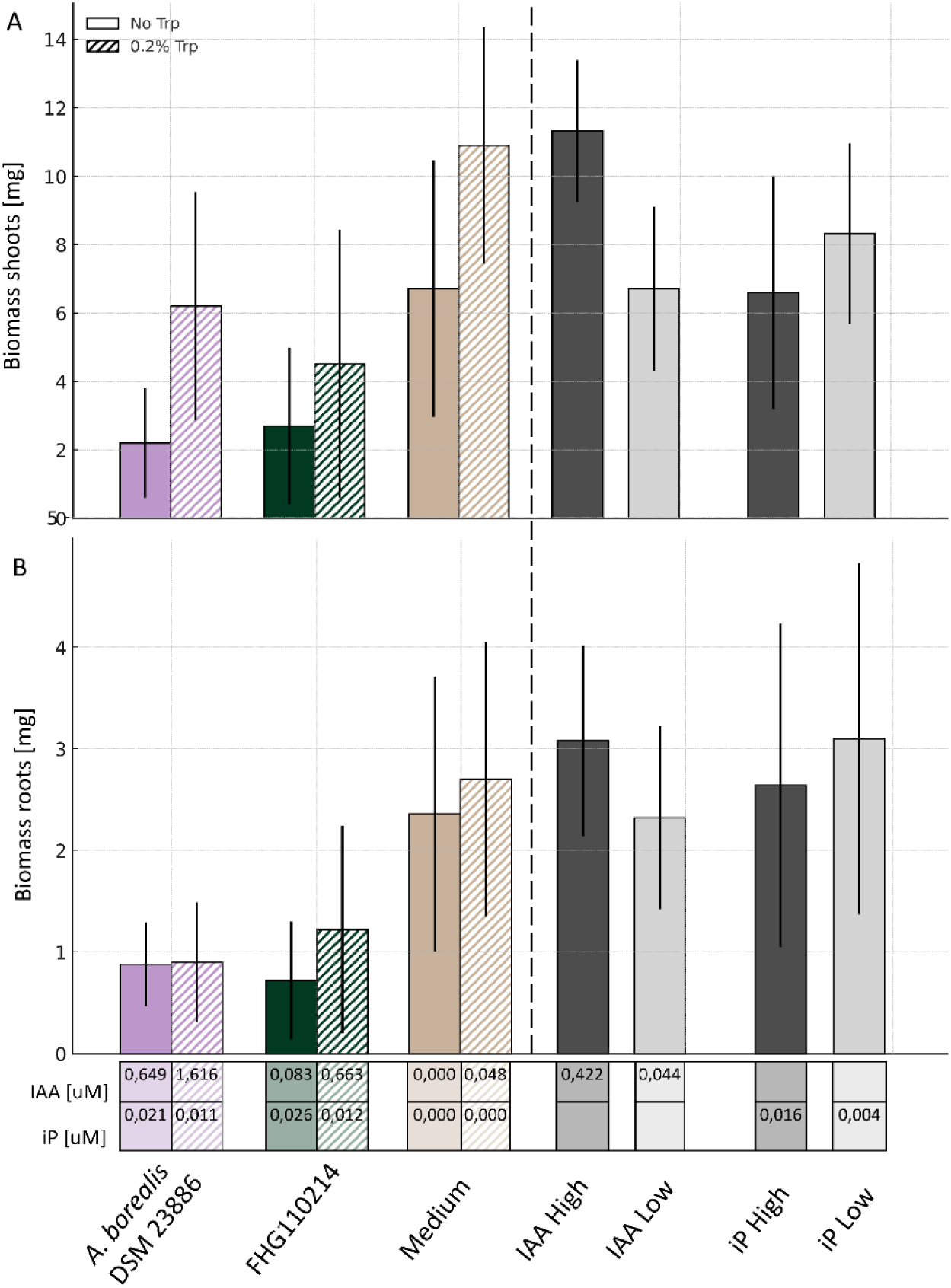
Dry Biomass of barley shoots (A) and roots (B) treated with extracts of strains *A. borealis* DSM 23886 (purple), FHG 110214 (green). Striped bars indicate extracts from Trp-supplemented cultivations, while solid bars indicate that extracts were not supplemented with Trp. Media controls (brown), as well as IAA and iP as pure compounds(grey), were also tested. The table indicates the final concentrations of phytohormone applied to the barley seedlings. In general, supplementation with Trp results in increased biomass, especially in the shoots. While pure compound controls show comparable growth to the media control, the strain extracts inhibit plant growth. This negative effect i s diminished upon supplementation with Trp, especially in the shoots.

Interestingly, our findings, along with those of Kielak *et al*., (31) and Gonçalves *et al*. (20), suggest that phytohormone production is common among Acidobacteriota, particularly among strains isolated from decaying wood or lignocellulosic substrates. This raises intriguing questions about the ecological roles of Acidobacteriota strains producing compounds like IAA and iP in natural systems (57).

### Result 4 – In-Depth Bioinformatic Analysis of PGP Traits in Acidobacteriota and FHG *Acidobacteriaceae*

Taken together, these results confirm that *Acidobacteriaceae* produce phytohormones in relevant amounts, but their role in PGP remains unresolved. Because iP can arise from the degradation of prenylated tRNAs in many bacteria (58), we asked whether *Acidobacteriaceae* also encode dedicated cytokinin pathways (involving lonely guy (LOG) enzymes) and whether these co-occur with additional PGPTs. Furthermore, we were interested in the biosynthetic mechanisms responsible for producing phytohormones. For batch analysis of the 618 genomes previously analysed, we employed a DIAMOND workflow against the PLABASE database (see materials and methods) and summarised hits as coverage (% of PGPT families detected). For the analysis of taxonomic patterns, we aggregated the results by GTDB-Tk family status, only considering acidobacterial families that have been scientifically described or have more than 20 representative genomes available (n = 391, 64.2% of the total dataset). Figure 5 represents the percentage coverage of PGPTs categorised by LABA at “le el two”. Values were hierarchically clustered to visualise similarity between different families and PGPTs. The analysis separated the families into two distinct clusters.

**Figure 5:**
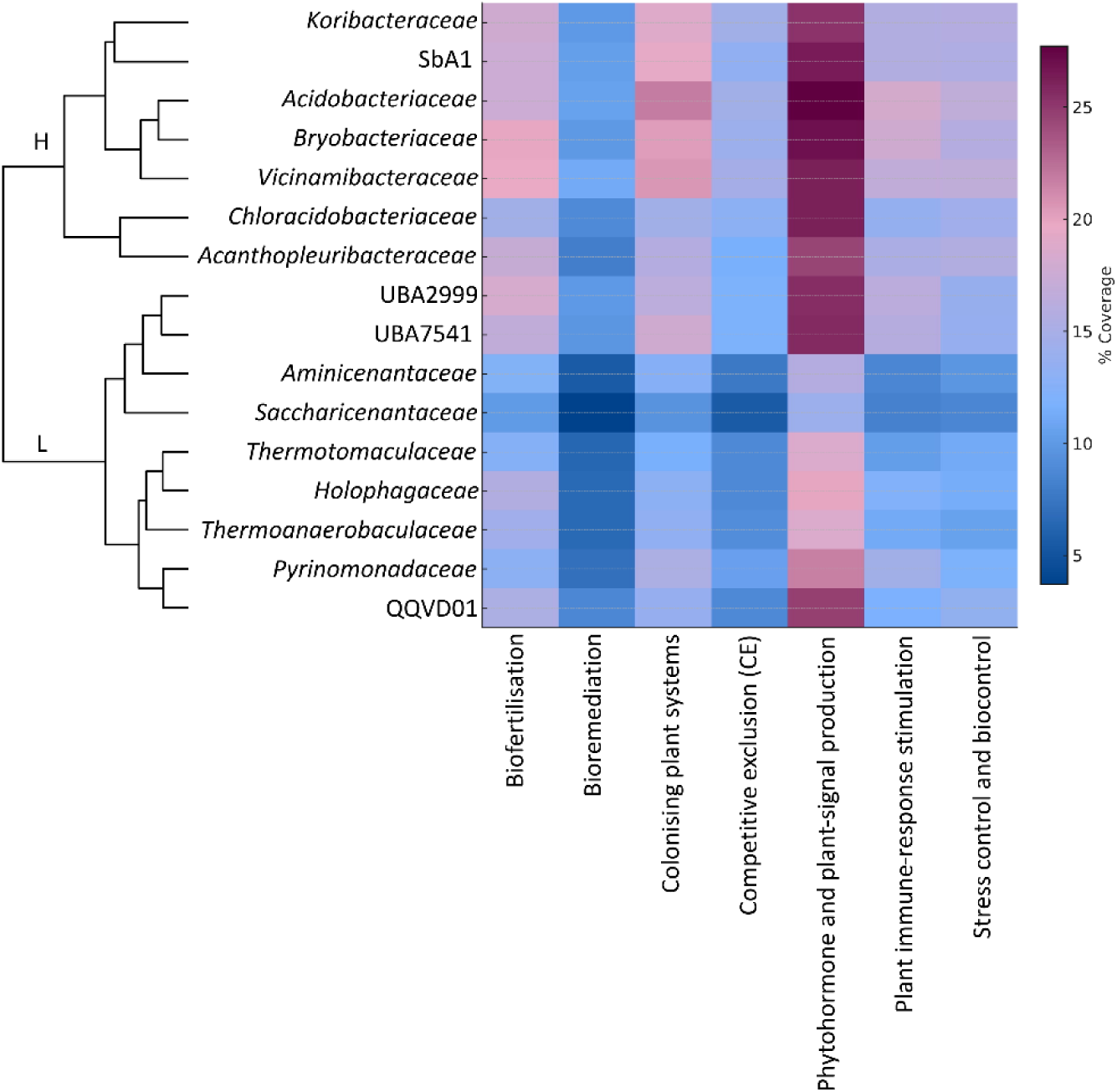
PGPT coverage across Acidobacteriota families. The heatmap displays the mean % coverage of PLABASE PGPT categories (level 2) for families with formal names or more than 20 genomes (n = 16). Coverage values were calculated as the proportion of detected PGPT families per functional category, relative to the total number of families defined in the PLABASE ontology. Rows and columns are hierarchically clustered. Two distinct groups emerge, indicating lower (L, bottom) and higher (H, top) PGPT coverage. The latter exhibits higher potential, particularly in the production of phytohormones and plant signals.

The nine families in the lower-coverage cluster (L) (UBA2999, UBA7541, *Aminicenantaceae*, *Saccharicenantaceae*, *Thermotomaculaceae*, *Holophagaceae*, *Thermoanaerobaculaceae*, *Pyrinomonadaceae*) show a comparatively low average PGPT potential (12.7±1.2%) with uniformly modest values across categories such as biofertilisation: 14.3 ± 1.4%; colonising plant system: 13.7 ± 1.4%; stress-control-biocontrol: 11.6 ± 1.0% and especially low bioremediation: 7.2 ± 0.9%. In contrast, families in the higher-coverage cluster H (*Koribacteraceae*, SbA1, *Acidobacteriaceae*, *Bryobacteraceae*, *Vicinamibacteraceae*, *Chloracidobacteriaceae*, *Acanthopleuribacteraceae*) showed consistently higher coverage with representative means of 26.1 ± 2.6% for plant signal production, 18.8 ± 2.4% for colonising plant system and 13.7 ± 1.7% competitive exclusion (overall range ∼17–26% across categories. Across both clusters, the bioremediation category is underrepresented, while the phytohormone and plant signalling group ranks top among both clusters. Overall, the families of *Bryobacteriaceae*, *Acidobacteriaceae*, and *Vicinamibacteriaceae* exhibit the highest levels of PGPTs. Notably, the *Acidobacteriaceae* perform best in the colonising plant system group (e.g., covering EPS production) and the phytohormone/plant signal production category.

We analysed DIAMOND outputs for enzymatic equipment supporting IAA production in our FHG *Acidobacteriaceae* strains. In bacteria, three pathways are described (50): indole-3-pyruvic acid (IPyA), indole-3-acetamide (IAM) and tryptamine (TAM). In the IPyA route, Trp is transaminated to indole-3-pyruvate, decarboxylated to indole-3-acetaldehyde, and oxidised to IAA; the IAM pathway converts Trp to indole-3-acetamide, then hydrolyses it to IAA; the TAM pathway decarboxylates Trp to tryptamine, which is subsequently oxidised via intermediates such as indole-3-acetaldehyde to IAA. All FHG *Acidobacteriaceae* include the key enzymes to synthesise Trp (trpA-E) and terminal enzymes for IAA production, such as an *amiE* homolog for the hydrolysis of indole-3-acetamide, and two aldehyde-dehydrogenases that may catalyse the final step of IAA production in both the TAM and the IPyA pathways. However, the automatic setup did not detect intermediate enzymes. Metabolomics using reference standards identified indole-pyruvic acid (CAS: 392-12-1) upon Trp supplementation, whereas tryptamine (CAS: 61-54-1) was not detected in FHG110202, FHG110214 and *A. borealis* DSM23886. FHG110511 showed neither intermediate. These observations support IPyA as the predominant route over TAM, while IAM remains a possible but unconfirmed option.

The production of iP is mediated by two distinct pathways; the t-RNA isopentenyltransferase (IPT) pathway involves the isoprenylation of tRNAs by t-RNA IPTs (*miaA*) and the context-dependent generation of iP through the degradation of these tRNAs (59, 60). The pathway is found ubiquitously in bacteria. Instead, the second pathway is less common and is directly associated with regulated cytokinin output in bacteria, which usually interact with plants. Here, the adenylate IPT modifies adenosine nucleotides using dimethylallyldiphosphate (61), thereby producing iP-like nucleotides further activated by lonely guy (LOG) enzymes (62, 63). Our genomic survey revealed that FHG110511, as well as FHG110214 and *A. borealis* DSM 23886, contain the tRNA-IPT (*miaA*) and accessory genes, while FHG110202 additionally contains a LOG (lonely guy) enzyme. Instead, the adenylate IPT could not be detected in any of the FHG *Acidobacteriaceae*. As indicated by the literature, the *miaA* gene is omnipresent in our acidobacterial dataset, while the LOG enzymes are less present (477/618) but still highly abundant in various acidobacterial classes. Notably, the Trp-responsive iP production aligned with our pathway content. In the strains harbouring the tRNA IDT pathway alone, we determined a reduced iP production upon Trp supplementation. This might be explained by an overall decrease in tRNA turnover due to the high availability of Trp, resulting in a subsequent reduction in global translation and, consequently, lower cytokine release through tRNA degradation mechanisms. Instead, FHG110202, the only inspected strain that harbours a LOG enzyme, does not show a Trp-dependent decrease in iP production, thus possibly less prone to global protein biosynthesis and tRNA turnover rates.

We also specifically looked for the presence of vital PGPT factors that could be tested in wet lab experiments. This included plate assays indicating phosphate solubilisation, nitrogen fixation and ACC-deaminase activity. The enzyme ACC deaminase (AcdS) breaks down 1-aminocyclopropane-1-carboxylic acid into ethylene, thereby playing a vital role in the plant defence mechanism. We did not detect *acdS* homologs in our FHG *Acidobacteriaceae* strains. Consistently, we could not detect any growth on plates containing 1-aminocyclopropane-1-carboxylic acid (data not shown). The enzyme AcdS, correlated to PGPT family number 7345, was also searched for in the entire dataset, but was not detected in any strain. The ability to solubilise phosphate from phytate was evaluated on medium containing phytate (data not shown). However, even though we did not detect any growth on agar plates, we identified genes related to phosphate solubilisation through genome mining. *A. borealis* DSM 23886(1.02e-119), FHG110214 (4.91e-107), and FHG110511 (1.57e-148) contained homologs of *appA* (PGPT2560), a phytase known to release Pi from organic phosphorus via a phytase or nucleotidase reaction. We also searched for phytase homologs belonging to PGPT families 2560, 2580, and 2585 in the acidobacterial dataset. Phytases were present in 117 genomes, but only in the classes of *Acidobacteriae* and *Blastocatellia*. The ability to fix nitrogen has been described for the Acidobacteriota before. Recently, Gonçalves *et al.* (20) described the presence of nif clusters in members of the *Holophagaceae* and *Acidobacteriaceae* families. In our genome analysis, we confirmed that members of the *Holophagaceae (Holophagaceae bacterium isolate CTSoil_007)* and *Acidobacteriaceae* (Acidobacteria bacterium isolate SZAS TMP-7) contained nitrogenase clusters encompassing all required *nif* genes A-F (PGPT families PGPT1-30). Besides the *Holophagaceae*, *nif* genes were also detected in *Bryobacteraceae* TMP-7, but not in any of the FHG *Acidobacteriaceae*.

Compared to the large-scale metagenomic analysis of Gonçalves *et al.*, our PGPT analysis shows both shared and divergent trends across the acidobacterial families. Some families, such as *Acidobacteriaceae*, *Thermoanaerobaculum*, *Aminicenantesaceae*, *Holophagaceae*, and *Koribacteraceae,* are grouped similarly in terms of their overall equipment, either uniformly high or low; other families show significant discrepancies. Specifically, our results diverge for the families *Bryobacteriaceae, Vicinamibacteraceae, Pyrinomonadaceae, and Chloracidobacteriaceae,* where the PGP potential differs substantially. These differences likely arise from the limited overlap in representative genomes between the two datasets. Overall, 9.2% of our dataset is also covered by the Goncalves dataset. Additionally, our analysis includes seven additional Acidobacteriota families. Notably, the dataset coverage is particularly low for most families with detected discrepancies, including *Bryobacteriaceae* (1/74), *Vicinamibacteraceae* (2/6), and *Pyrinomonadaceae* (2/87). The overlap of the *Chloracidobacteriaceae* (10/14) is relatively high; however, in addition to the dataset composition, methodological differences likely also contribute to the differences. Gonçalves *et al*. focused on 91 curated PGPTs, whereas our genome mining approach utilised the complete PLABASE database, encompassing over 1,300 plant growth-promoting gene families. This broader functional scope may capture additional trait diversity and help explain the differential PGPT evaluation and distribution observed across families.

The PGPT profiles of the FHG *Acidobacteriaceae* are consistent with both our broader family-level analysis (Figure 5) and the patterns reported in the large-scale metagenomic study by Goncalves *et al*. (2024). In all cases, *Acidobacteriaceae* show a high frequency of PGPTs, indicating a conserved functional repertoire within this lineage. Notably, Gonçalves *et al*. also identified a similar PGPT pattern in this family, including genes associated with phytohormone biosynthesis and phytase activity. These congruent findings suggest that *Acidobacteriaceae* may play a robust and stable role in plant–microbe interactions across diverse environments.

### Result 5 – Effects of Tryptophan on Metabolite Production and Phytopathogen Inhibition

Trp is abundant in the root exudates of many plants, and is described to be used by bacteria to synthesise auxins (64). Indeed, the synthesis of phytohormones by rhizosphere bacteria often depends on the presence of Trp precursors in the root exudates (65). Thus, this metabolic shift might reflect an environmental response of the strains towards the Trp-enriched plant environment (66). To investigate whether supplementation with Trp induces a global metabolic shift or stimulates the production of phytohormones, and to potentially describe the regulated metabolites, we performed time-resolved untargeted metabolomics analysis. Extracts cultivated with and without Trp were analysed with ultra-high-performance liquid chromatography–quadrupole time-of-flight high-resolution mass spectrometry (UHPLC-QTOF-HRMS), and the calculated feature table was filtered to remove features identified as noise. Subsequent principal component analysis (PCA) of log2-transformed feature intensities (Figure S14) revealed that the samples did not exhibit a strict separation by strain or cultivation day, but instead were separated by supplementation status, indicating a broad metabolomic shift associated with Trp supplementation. Furthermore, we observed temporal metabolic shifts that were more distinct for *A. borealis* DSM 23886, especially if supplemented with Trp.

To further investigate Trp-dependent metabolomic responses, we filtered and categorised features based on their presence-absence and fold-change behaviour in response to Trp supplementation. Briefly, features were retained if they were present in both bucketing calculations. Following filtering, feature intensities were compared between the uninduced and induced conditions to derive fold changes. Based on those, we assigned biological categories from “extinguished” to “switched-on”. Individual features were then plotted and coloured to indicate their biological response (Figure S15). The Barcoding plot confirmed the metabolic shift in response to Trp supplementation. We observed differences in media consumption (left side) and changes in features that were clearly strain-dependent (right side). Overall, there is an apparent change in the metabolism indicated by the number of features that change upon Trp supplementation. For *A. borealis* DSM 23886, 165 features are detected on day 14 without Trp, while 684 features (4.1x) are present if Trp is supplemented. The same trend is observed in FHG110214. Here, the feature number increases from 109 (without Trp supplementation) to 337 (3.1x) (with Trp supplementation). We independently verified this result in a second cultivation (Figure S15, bottom half). Here, a similar fold change is observed. While the feature number increases 3.2-fold in *A. borealis* DSM 23886, it exhibits a similar behaviour in FHG110214 (2.8-fold). Those numbers already indicate a strong metabolic activation. This is supported by the fact that, considering all features present under both conditions, 80% of the features are at least lightly upregulated, while 19% are downregulated, and only 1% remain stable.

To further investigate Trp-dependent metabolomic responses considering media and strain-specific features, we classified Trp-activated uptake as referring to features initially present in the medium that were consumed only in Trp-induced cultures. In contrast, Trp-inhibited uptake describes the opposite: features consumed only in uninduced cultures. The rest of the classifications remained the same. Features that were stable in the medium and did not change in intensity were removed from the analysis. Figure 6 (top bar chart) displays the percentage distribution of features across these categories. The “switched on” class accounts for the largest fraction in both strains (*A. borealis* DSM 23886: 56.8%; FHG110214: 45.1%). Specifically, 10.4 and 16.9% of features exhibit an “increased production” profile while 7.5 and 6.3%, respectively, are “switched off.” mall proportions (below 10%) represent different metabolisation patterns of media components under induced and uninduced conditions.

**Figure 6:**
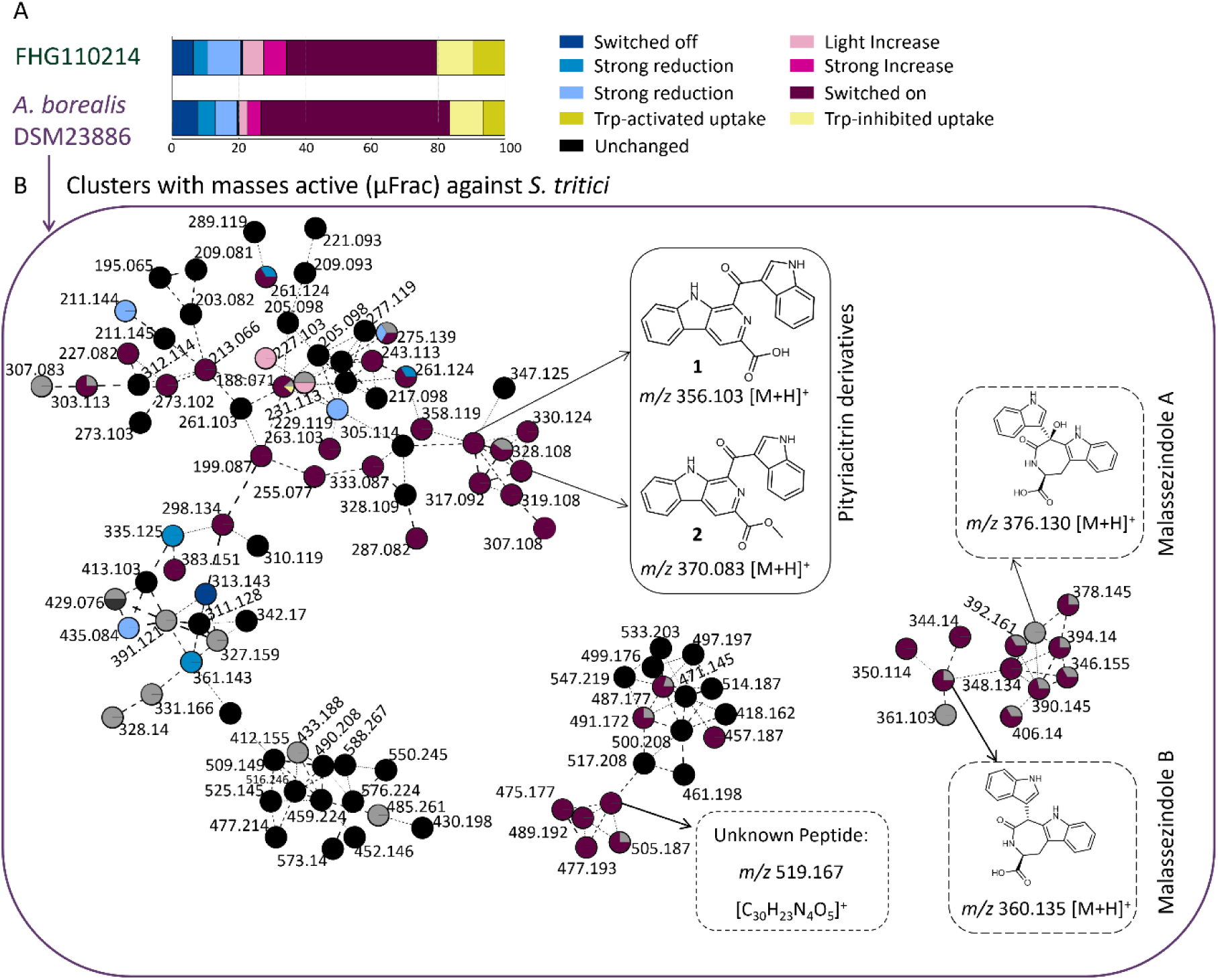
Overall metabolic response of cultivation with Trp in strains FHG 110214 and *A. borealis* DSM 23886. (A). Molecular network of masses active against *S. tritici* MUCL 45407 as detected by µ Frac-coupled screening of *A. borealis* DSM 23886 extracts (B). (A) After feature detection, the relative presence of masses with and without Trp was compared for categorisation. Categories were defined as follows: “ switched - off” (dar - blue, no feature detected upon Trp supplementation) and “ switched - on” (dar - red, Feature only present upon Trp supplementation). Media components that were metabolised differently upon Trp supplementation are shown in yellow. (B)Molecular Networking of MS/ MS data generated from a Trp-induced *A. borealis* DSM 23886 culture, limited to clusters containing masses that showed antifungal activity against *S. tritici* MUCL 45407. Active masses were all specifically “ switched on” when supplemented with Trp. While some of them remained unknown (e. g. 519.167 m/ z peptide), others were dereplicated by comparison with database standards (malassezindole A, B) or isolated and structure elucidated(pityriacitrin B and pityriacitrin B methyl ester).

Given the pronounced metabolic changes observed in Trp-induced extracts, we were encouraged to re-enter our initial bioactivity discovery approach, now particularly testing the generated extracts from FHG110214 and *A. borealis* DSM 23886 for activity against the phytopathogenic fungus *Septoria tritici* MUCL45407, a residue-borne wheat pathogen causing *Septoria tritici* blotch (STB), one of the most significant diseases of wheat (67). By testing our extracts in a microbroth dilution assay, we indeed determined a strong growth-inhibitory effect, exclusively in the samples that underwent Trp supplementation. For the identification of activity-causing compounds, one representative active extract of FHG110214 and *A. borealis* DSM 23886 was microfractionated and re-screened against *S.tritici* MUCL 45407. Based on the molecular formula and the observed MS/MS fragmentation pattern, several Trp-related compounds were dereplicated within the active fractions of both strains, with a confidence level 2 (putatively annotated metabolites (44)). This enabled the annotation of hydroxyglyantrypine (68) as well as malassezindoles A, B (69) and putative yet undescribed derivatives thereof (m/z: 148.1349, 376.1293, 360.1348, 362.1498). Notably, malassezindoles production is known to be strongly dependent on Trp as the sole nitrogen source in their fungal production strains (70). This observation aligns well with our cultivation conditions and experimental setup. However, since the malassezindoles have not been described for bacteria yet, we have expanded our quality control by including regular agar plating, which is conducted concurrently with sampling at the beginning and end of cultivation. Additionally, we continued to perform 16S/18S sequencing from liquid culture samples. Agar plates were incubated for at least 14 days, documented, and routinely inspected using a binocular microscope. Neither measure raised any concerns about handling fungal-contaminated samples. Besides malassezindoles, we annotated the β-carbolines acetyl-beta-carboline, pityriacitrin B (69, 71, 72) and a methylated derivative thereof. Pityriacitrins are produced by fungi and bacteria (73) and recognised for their UV-protective properties (74). Furthermore, the β-carbolines scaffold has been recently reviewed in the context of antifungal activity (75). Indeed, pityriacitrin A, B, and their methyl esters have been described to inhibit fungal growth (73, 76). Furthermore, pityriacitrin B was shown to have protective activity against fungal colonisation of plant hosts (73). In bacterial systems, beta-carboline cores, such as those found in pityriacitrin, are formed via Pictet-Spenglerase enzymes (77–79). Instead, biosynthesis of the Trp-dependent metabolites, closely related to the malassezindoles, has been described for the smut fungus *Ustilago maydis*, where a single enzymatic reaction, catalysed by the enzyme Tam1, is essential for their formation, converting Trp to indolepyruvate (80). Although the biosynthetic pathways for these compounds have not been elucidated in bacteria, a similar mechanism can be inferred. However, a BLAST search for analogous proteins within our acidobacterial genome dataset did not yield any significant hits. An additional signal observed in the active fraction, co-eluting with malassezindoles under the applied chromatographic conditions, suggested the presence of a peptide, with an m/z value of 519.1655 [M+H]+ corresponding to the molecular formula C_30_H_22_N_4_O_5_.

We focused our isolation efforts on compounds previously reported to exhibit antifungal properties. Preparative and semi-preparative isolation methods were employed on the fractions containing malassezindoles and their derivatives. Two compounds were purified, and their structures were elucidated by NMR analysis in DMSO-*d*_6_, employing ^1^H, ^13^C, COSY, HSQC, HMBC, TOCSY, ROESY and NOESY measurements (Table S3-4). Compound **1** (C₂₁ ₁₃N₃O₃) was identified as pityriacitrin B while compound **2** (C₂₂ ₁₅N₃O₃) was determined to be its methyl ester. The spectroscopic data for both compounds were consistent with literature reports (72, 81). Detailed ^1^H and ^13^C NMR data, as well as key COSY, ROESY and HMBC correlations, are summarised in the Supporting Information (Tables S2–S3).To examine the features previously identified to be bioactive, we mapped them onto our categorised feature dataset. This allowed us to assess which metabolic response classes these metabolites belong to. Additionally, we explored other Trp-dependent masses by molecular networking, aiming to identify novel or derivative compounds using database annotations.

The observed changes in activity profiles and the Trp-dependent modulation of extracts underscore the impact of Trp supplementation on the production of specific, bioactivity-causing compounds. Having already linked several of these induced features to bioactivity through microfractionation, we next turned to molecular networking to explore their structural context within the overall metabolome. This approach enabled us to visualise whether the bioactive compounds belong to larger molecular families, identify structurally related analogues, and determine whether induction leads to the coordinated activation of specific metabolic clusters. By integrating activity data with MS/MS-based network topology, we aimed to prioritise entire subnetworks of interest and assess how broadly Trp supplementation influences distinct chemical families across strains.

Acidobacteriota often dominate soil bacterial communities, particularly in acidic and oligotrophic environments, yet their functional roles remain underexplored. Our findings contribute to growing evidence that members of this phylum possess underestimated biosynthetic and ecological potential, including the ability to produce phytohormones and possibly bioactive metabolites with the potential to antagonise fungal growth.

Beyond our cultivated FHG *Acidobacteriaceae*, genome-mining of 618 high-quality Acidobacteriota genomes offers a complementary perspective to the large-scale MAG-based analysis by Gonçalves *et al*. (2024), which analysed 758 MAGs. Both studies identify *Acidobacteriaceae*, *Bryobacteriaceae*, and *Koribacteraceae* as families with enriched PGPT potential, while families such as *Holophagaceae* and *Vicinamibacteraceae* appear comparatively under-equipped. However, notable discrepancies emerged, likely reflecting differences in the underlying datasets and the methodological approach.

Despite the current technical limitations, exploring Acidobacteriota as sources of novel bioactive compounds or as contributors to plant–soil health remains a promising avenue. A considerable limitation, however, is their challenging growth in laboratory settings. If obtained as isolates, their total cell biomass remained low, even when using optimised media conditions, and some strains even show aggregated growth forms. Given the intriguing biosynthetic potential predicted from the genome, more efforts are needed to characterise and exploit this phylum, including, for example, the development of genetic tools for in situ activation, overexpression, and deletion of specific genes, as well as the establishment of a genetically tractable heterologous expression system in Acidobacteriota-hosts. Together, this could lead to a better understanding and utilisation of Acidobacteriota, aiming to discover new methods for their role in sustainable agriculture, bioremediation, and natural product discovery.

## Methods

### Microbial cultivation of acidobacterial strains

Acidobacterial strains were routinely cultivated in R2A Medium (R2A Broth, HIMEDIA, Maharashtra, India) buffered with MES-hydrate(2-(N-Morpholino)ethanesulfonic acid hydrate, CAS: 1266615-59-1) (1.95 g/l) at pH 6.0. Cultures were incubated at 28°C with 75 rpm. To sustain healthy and continuous growth, 10% of the 7-day-old preculture (30 ml in 100 ml Erlenmeyer flask) was used to inoculate the main cultures (100 ml in 300 ml Erlenmeyer flasks). For quality control, we transferred 100 μl after each inoculation or sampling step to an R2A-MES-buffered agar plate to observe colony morphologies and identify the growth of contaminant microorganisms. Additionally, we performed 16S rRNA gene amplification testing with primer pair E8F (50-GAG TTT GAT CCT GGC TCA G-30) and 1492R (50-ACG GYT ACC TTG TTA CGA CTT-30) (Lane, 1991). For the OSMAC cultivation approach, we performed 24-well-scale and Erlenmeyer flask-scale cultivation. We prepared forty 24-well Duetz system plates with 20 different media (each medium 4 ml) and autoclaved the plates before inoculation (10%) with a preculture grown for 14 days. Applied media were diluted medium 5254 (82)(2 g soluble starch, 2 g glycerol, 0.5 g corn steep liquor 1 g peptone, 0.4 g yeast extract, 0.2 g NaCl; pH 5.5, buffered with 1.95 g/L MES), HD (0.5 g casein peptone, 0.1 g glucose, 0.25 g yeast extract; pH 5.0, unbuffered; DSMZ ID 1135), and PSYA 5 (26)(1.8 g KH_2_PO_4_, 0.2 g MgSO_4_ x 7H_2_O, 1 g yeast extract; optionally with addition of sucrose at 0.5% or 3%; pH 5.0, unbuffered) as well as a diversity of VL55-based and R2A-based (1-fold, 2-fold, 4-fold) media generated by optional addition of different carbon sources (Pektin, GlcNAc – 0.05%; Xylan, Cellobiose – 0.5%; and Sucrose 3%; applied to 2-fold R2A)(Table S2) (25, 39). Furthermore, as a surface attachment enabling strategy, we added ceramic beads (hollow cylinders; 3 pcs/well plus 4 ml of media), quartz beads (round; 10 pcs/well plus 4 ml of media), or solidified the respective medium using 0.8% phytagel. The plates were incubated at 28°C, shaking at 120 rpm or with manual shaking once per day. Cultivation in a 300 ml Erlenmeyer flask was executed in 100 mL media by inoculation (10%) with a preculture grown for 7 days. The cultures were incubated at 28°C for 14 days with or without shaking at 75 rpm. We added 20 g of ceramic beads (Eheim mech) to non-shaking cultures. After cultivation, cultures were frozen, freeze-dried and extracted (see below).

For Trp-induced kinetic studies, we used a 1.2-fold concentrated R2A medium buffered with 2-(*N*-morpholino)ethanesulfonic acid (MES) at a pH of 6.0, which was supplemented in a 4:1 ratio with 1% sterile-filtered Trp (CAS 73-22-3) to a final concentration of 0.2% Trp and a 1-fold concentrated medium. Kinetic studies were performed in 300 ml flasks at least in duplicates. For sample collection, we transferred 4 ml of culture to 24-deep-well plates for further processing. In parallel, the remaining culture sample was used for quality control by agar plating (R2A-MES, pH 6.0) and 16S rDNA PCR. After the final harvest day, the 24-well plates were freeze-dried and extracted using methanol (SI-extended methods and results).

### Bioassays

Antimicrobial Screenings of crude extracts were performed using microplate broth dilution assays at the final assay concentration of 0.25-fold, 0.5-fold, one-fold and two-fold (in relation to the culture volume) against bacteria and fungi (*Escherichia coli* ATCC 35218, *Bacillus subtilis* DSM 10, *Mycobacterium smegmatis* ATCC 607, *Candida albicans* FH 2173 and *Septoria tritici* MUCL 45407) as described previously (39). Crude extracts showing at least 70% growth inhibition were considered bioactive and were subjected to microfractionation for dereplication of bioactivity-causing metabolites (39). Testing of crude extracts on barley seedlings was performed in glass vials filled with 5 ml of SH medium, inoculated with one surface-disinfected barley seed per vial (SI-extended methods and results).

#### Creation and Curation of the Acidobacteriota Genome Dataset

As of March 2023, Acidobacteriota assemblies matching our selection criteria (NCBI:txid, filter: RefSeq annotation) were retrieved from NCBI in .gbff file format. In addition, four Acidobacteriota assemblies, from strains isolated from termite nest material (FHG110202, FHG110214), soil (FHG110511) and a genome generated for *A. borealis* DSM 23886 complemented the dataset (n = 1425). Genome assemblies were selected based on their completeness and contamination using CheckM2(40). Only high-quality genome assemblies defined by MIMAG (41) evaluation criteria demonstrating 90-100% completeness and 0-5% contamination were retained for further evaluation (n = 648). For this, redundancies and taxonomic affiliation were checked by GTDB-Tk (42). In total, 618 curated genome assemblies were processed for metadata evaluation, BGC potential, and PGPT mining (SI-extended methods and results for details). To correlate the genomic data to the taxonomic affiliation, a whole-genome sequence-based phylogenetic tree of the GTDB-generated multiple-sequence alignment file was visualised using iTOL v4 (45).

#### Genomic Analysis of PGPT Traits

The coding sequences from the previously described genomic dataset were extracted using a custom Python pipeline built with BioPython (SeqIO). To generate FASTA documents for further processing, we retrieved the gene name, product description and translated protein sequence for each CDS. For pre-existing FASTA files, the extraction of CDSs can be skipped. To document the number of CDSs extracted per genome, a log file was generated. The PLABASE PGPT database (83) (PGPT_BASE_nr_Aug2021n_ul_1.dmnd) was used as a reference for a similarity search against the extracted protein sequences using DIAMOND v2. The script supports various sensitivity modes, enabling users to select the most suitable mode based on their specific input. The parameters and applied evaluation thresholds used in this study are described in the Extended Methods and Results section of the supplementary information.

#### Mass spectrometry

Ultra-high-performance liquid chromatography (UHPLC) coupled with quadrupole time-of-flight (QTOF) high-resolution mass spectrometry (HR-MS) and tandem MS (MS/MS) was performed using an Agilent 1290 Infinity LC system connected to a maXis II QTOF mass spectrometer (Bruker Daltonics) equipped with an electrospray ionisation (ESI) source. Separation of samples was achieved on an Acquity UPLC BEH C18 column (130 Å pore size, 1.7 µm particle size, 2.1 × 100 mm) fitted with a matching pre-column (2.1 × 5 mm, 1.7 µm, 130 Å) at a temperature of 45°C. Chromatographic separation was achieved using a linear gradient elution of solvent A (water with 0.1% formic acid) and solvent B (acetonitrile with 0.1% formic acid) at a constant flow rate of 0.6 ml/min. The gradient program was as follows: 95% A at 0.00–0.30 min, gradually decreasing to 4.75% A by 18.00 min, then to 0% A at 18.10 min, held until 22.50 min, returned to 95% A at 22.60 min, and equilibrated until 25.00 min. Mass spectra were recorded in positive mode over a mass range of m/z 50–2000 with a spectra rate of 1 Hz. MS/MS acquisition was performed at a 6 Hz scan rate, targeting the top five most intense ions per full mass spectrum for collision-induced dissociation (CID) using nitrogen, as per the table published by Eichberg *et al.* (84). Precursors were excluded after two spectra, with a release duration of 0.5 min and re-e aluated if their signal intensity increased by ≥1.5-fold. All data analysis was performed with DataAnalysis 4.4 (Bruker, Billerica, MA, USA).

#### Chemotype-barcoding matrix, Multivariate analysis and statistics

Raw data processing was performed using DataAnalysis 5.2 (Bruker) as previously published, and the complete data set was copied and processed with both line spectra thresholds (here: 5,000 and 10,000) (85). Bucketing was performed on both sets at the same time, resulting in one table containing all buckets deemed identical for every sample (under both thresholds) and by applying the data curation measures as described in the extended methods and results of the supplementary information. This analysis revealed a final table of 219 samples with 3,092 buckets (OSMAC) and 48 samples with 2,759 buckets (Trp-Induction), which were used for the respective barcode generation and for categorising. The generated bucket tables were used as (i) a binary presence/absence matrix (result 1) or (ii) a quantitative intensity matrix (result 5). For PCoA on the binary matrix, (i) Jaccard dissimilarities were computed (skbio.diversity.beta_diversity, metric=“jaccard”), and PCoA (skbio.stats.ordination.pcoa) was done using scikit-bio (v0.7.0). PERMANOVA (skbio.stats.distance.permanova) assessed group differences with 999 permutations. We tested the effects of medium, cultivation size (small-scale vs. large-scale), and bead condition (Quartz/Ceramic/None) in a small-scale subset. For PCA on quantitative data (ii), we log2-transformed peak area intensities using scikit-learn (v1.3) in Python. Before PCA, undetected features were defined as zero, and samples were normalised to the injection volume wherever indicated. All visualisations were generated using Matplotlib, with attributes changed based on metadata information through shape, colour, and border style to reflect replicates and experimental conditions.

### Compound Isolation

The crude extract was dissolved in 10% MeOH to perform SPE utilising a 2 L column packed with Amberlite XAD-16 N and step gradient elution of 10%, 20%, 40%, 60%, 80%, and 100% MeOH. Based on LC-MS analysis, several fractions containing the compounds of interest were combined and evaporated to dryness. During isolation, semi-preparative and preparative HPLC were performed on Agilent 1100 HPLC systems equipped with external Gilson fraction collectors using H2O + 0.1% FA and ACN + 0.1% FA as mobile phases. Final purification was achieved by UPLC fractionation using the chromatographic method described in the UPLC-MS section, with fraction collection of 16 seconds per fraction, starting directly after injection.

### Structure elucidation by NMR

NMR spectra were recorded on a Bruker AVANCE III HD 700 MHz spectrometer operating at ^1^H frequency of 700 MHz and a ^13^C-carbon frequency of 140 MHz. The instrument was equipped with a 5 mm TCI cryo probe head. COSY, TOCSY, ROESY, NOESY, HSQC, and HMBC experiments were recorded using standard pulse programs. All isolated samples were measured in DMSO-d6. Chemical shifts were referenced to the solvent signals (^1^H: 2.50 ppm, ^13^C: 39.5 ppm)

## Supporting information

SummaryAcidobacteriota

SI_ExploringAcidobacteriota

## Acknowledgements

We would like to thank Oliver Schwengers for the assembly and annotation of the genomes. Prof. Dr. Sylvia Schnell and Rita Geißler-Plaum for scientific and technical support of the plant-related experiments. Walter Lanzalonga was supported by the Erasmus+ Programme of the European Union through a student mobility grant. Celine Zumkeller was supported by the dissertation completion grant, offered based on the Gender Equality Concept provided by Justus-Liebig-University Gießen.

## Abbreviations

OSMAC: one-strain–many-compounds
IAA: indole-3-acetic acid
iP: N6-(Δ - isopentenyl)adenine
PGPT: plant growth–promoting trait
BGC: biosynthetic gene cluster
NRP: nonribosomal peptide
PKS: polyketide synthase
RiPP: ribosomally synthesized and post-translationally modified peptide
MAG: metagenome-assembled genome
UMAP: Uniform Manifold Approximation and Projection
GTDB: Genome Taxonomy Database
antiSMASH: antibiotics & Secondary Metabolite Analysis Shell
iTOL: Interactive Tree Of Life
UHPLC-QTOF-HRMS: ultra-high-performance liquid chromatography–quadrupole time-of-flight high-resolution mass spectrometry
DMSO: dimethyl sulfoxide
MES: 2-(N-morpholino)ethanesulfonic acid
Trp: tryptophan

## References

1. Jones RT, Robeson MS, Lauber CL, Hamady M, Knight R, Fierer N. 2009. A comprehensive survey of soil acidobacterial diversity using pyrosequencing and clone library analyses. The ISME Journal 3:442–453.

2. Koch IH, Gich F, Dunfield PF, Overmann J. 2008. Edaphobacter modestus gen. nov., sp. nov., and Edaphobacter aggregans sp. nov., acidobacteria isolated from alpine and forest soils. International Journal of Systematic and Evolutionary Microbiology 58:1114–1122.

3. Eichorst SA, Breznak JA, Schmidt TM. 2007. Isolation and Characterization of Soil Bacteria That Define Terriglobus gen. nov., in the Phylum Acidobacteria. Appl Environ Microbiol 73:2708–2717.

4. Flieder M, Buongiorno J, Herbold CW, Hausmann B, Rattei T, Lloyd KG, Loy A, Wasmund K. 2021. Novel taxa of Acidobacteriota implicated in seafloor sulfur cycling. ISME J 1–22.

5. Mehta-Kolte MG, Bond DR. 2012. Geothrix fermentans Secretes Two Different Redox-Active Compounds To Utilize Electron Acceptors across a Wide Range of Redox Potentials. Appl Environ Microbiol 78:6987–6995.

6. Pankratov TA, Dedysh SN. 2010. Granulicella paludicola gen. nov., sp. nov., Granulicella pectinivorans sp. nov., Granulicella aggregans sp. nov. and Granulicella rosea sp. nov., acidophilic, polymer-degrading acidobacteria from Sphagnum peat bogs. International Journal of Systematic and Evolutionary Microbiology, 60:2951–2959.

7. Belova SE, Ravin NV, Pankratov TA, Rakitin AL, Ivanova AA, Beletsky AV, Mardanov AV, Sinninghe Damsté JS, Dedysh SN. 2018. Hydrolytic Capabilities as a Key to Environmental Success: Chitinolytic and Cellulolytic Acidobacteria From Acidic Sub-arctic Soils and Boreal Peatlands. Front Microbiol 9.

8. Sipes K, Buongiorno J, Steen AD, Abramov AA, Abuah C, Peters SL, Gianonne RJ, Hettich RL, Boike J, Garcia SL, Vishnivetskaya TA, Lloyd KG. 2024. Depth-specific distribution of bacterial MAGs in permafrost active layer in Ny Ålesund, Svalbard (79°N). Systematic and Applied Microbiology 47:126544.

9. Kishimoto N, Kosako Y, Tano T. 1991. Acidobacterium capsulatum gen. nov., sp. nov.: An acidophilic chemoorganotrophic bacterium containing menaquinone from acidic mineral environment. Current Microbiology 22:1–7.

10. Ludwig W, Bauer SH, Bauer M, Held I, Kirchhof G, Schulze R, Huber I, Spring S, Hartmann A, Schleifer KH. 1997. Detection and in situ identification of representatives of a widely distributed new bacterial phylum. FEMS Microbiology Letters 153:181–190.

11. Quaiser A, Ochsenreiter T, Lanz C, Schuster SC, Treusch AH, Eck J, Schleper C. 2003. Acidobacteria form a coherent but highly diverse group within the bacterial domain: evidence from environmental genomics. Molecular Microbiology 50:563–575.

12. Dedysh SN, Yilmaz P. 2018. Refining the taxonomic structure of the phylum Acidobacteria. International Journal of Systematic and Evolutionary Microbiology 68:3796–3806.

13. Ward NL, Challacombe JF, Janssen PH, Henrissat B, Coutinho PM, Wu M, Xie G, Haft DH, Sait M, Badger J, Barabote RD, Bradley B, Brettin TS, Brinkac LM, Bruce D, Creasy T, Daugherty SC, Davidsen TM, DeBoy RT, Detter JC, Dodson RJ, Durkin AS, Ganapathy A, Gwinn-Giglio M, Han CS, Khouri H, Kiss H, Kothari SP, Madupu R, Nelson KE, Nelson WC, Paulsen I, Penn K, Ren Q, Rosovitz MJ, Selengut JD, Shrivastava S, Sullivan SA, Tapia R, Thompson LS, Watkins KL, Yang Q, Yu C, Zafar N, Zhou L, Kuske CR. 2009. Three Genomes from the Phylum Acidobacteria Provide Insight into the Lifestyles of These Microorganisms in Soils. Appl Environ Microbiol 75:2046–2056.

14. Lee S-H, Ka J-O, Cho J-C. 2008. Members of the phylum Acidobacteria are dominant and metabolically active in rhizosphere soil. FEMS Microbiol Lett 285:263–269.

15. Dyksma S, Pester M. 2023. Oxygen respiration and polysaccharide degradation by a sulfate-reducing acidobacterium. Nat Commun 14:6337.

16. Foesel BU, Rohde M, Overmann J. 2013. Blastocatella fastidiosa gen. nov., sp. nov., isolated from semiarid savanna soil – The first described species of Acidobacteria subdivision 4. Systematic and Applied Microbiology 36:82–89.

17. Hausmann B, Pelikan C, Herbold CW, Köstlbacher S, Albertsen M, Eichorst SA, Rio TG del, Huemer M, Nielsen PH, Rattei T, Stingl U, Tringe SG, Trojan D, Wentrup C, Woebken D, Pester M, Loy A. 2018. Peatland Acidobacteria with a dissimilatory sulfur metabolism. ISME J 12:1729–1742.

18. Reji L, Zhang X. 2022. Genome-Resolved Metagenomics Informs the Functional Ecology of Uncultured Acidobacteria in Redox Oscillated Sphagnum Peat. mSystems 7:e00055–22.

19. Giguere AT, Eichorst SA, Meier DV, Herbold CW, Richter A, Greening C, Woebken D. 2021. Acidobacteria are active and abundant members of diverse atmospheric H2-oxidizing communities detected in temperate soils. ISME J 15:363–376.

20. Gonçalves OS, Fernandes AS, Tupy SM, Ferreira TG, Almeida LN, Creevey CJ, Santana MF. 2024. Insights into plant interactions and the biogeochemical role of the globally widespread Acidobacteriota phylum. Soil Biology and Biochemistry 192:109369.

21. Parsley LC, Linneman J, Goode AM, Becklund K, George I, Goodman RM, Lopanik NB, Liles MR. 2011. Polyketide synthase pathways identified from a metagenomic library are derived from soil Acidobacteria. FEMS Microbiology Ecology 78:176–187.

22. Crits-Christoph A, Diamond S, Butterfield CN, Thomas BC, Banfield JF. 2018. Novel soil bacteria possess diverse genes for secondary metabolite biosynthesis. Nature 558:440–444.

23. McReynolds E, Elshahed MS, Youssef NH. 2025. An ecological-evolutionary perspective on the genomic diversity and habitat preferences of the Acidobacteriota. Microbial Genomics 11:001344.

24. Leopold-Messer S, Chepkirui C, Mabesoone MFJ, Meyer J, Paoli L, Sunagawa S, Uria AR, Wakimoto T, Piel J. 2023. Animal-associated marine Acidobacteria with a rich natural-product repertoire. Chem 9:3696–3713.

25. Oberpaul M, Zumkeller CM, Culver T, Spohn M, Mihajlovic S, Leis B, Glaeser SP, Plarre R, McMahon DP, Hammann P, Schäberle TF, Glaeser J, Vilcinskas A. 2020. High-throughput cultivation for the selective isolation of Acidobacteria from termite nests. Front Microbiol 11.

26. Campanharo JC, Kielak AM, Castellane TCL, Kuramae EE, Lemos EG de M. 2016. Optimized medium culture for Acidobacteria subdivision 1 strains. FEMS Microbiology Letters 363:fnw245.

27. Sait M, Hugenholtz P, Janssen PH. 2002. Cultivation of globally distributed soil bacteria from phylogenetic lineages previously only detected in cultivation-independent surveys. Environmental Microbiology 4:654–666.

28. Kuske CR, Barns SM, Busch JD. 1997. Diverse uncultivated bacterial groups from soils of the arid southwestern United States that are present in many geographic regions. Applied and Environmental Microbiology 63:3614–3621.

29. Felske A, De Vos WM, Akkermans ADL. 2000. Spatial distribution of 16S rRNA levels from uncultured acidobacteria in soil. Lett Appl Microbiol 31:118–122.

30. Eichorst SA, Kuske CR, Schmidt TM. 2011. Influence of Plant Polymers on the Distribution and Cultivation of Bacteria in the Phylum Acidobacteria. Applied and Environmental Microbiology 77:586–596.

31. Kielak AM, Cipriano MAP, Kuramae EE. 2016. Acidobacteria strains from subdivision 1 act as plant growth-promoting bacteria. Arch Microbiol 198:987–993.

32. Damsté JSS, Rijpstra WIC, Dedysh SN, Foesel BU, Villanueva L. 2017. Pheno- and Genotyping of Hopanoid Production in Acidobacteria. Front Microbiol 8.

33. Kielak AM, Castellane TCL, Campanharo JC, Colnago LA, Costa OYA, Corradi da Silva ML, van Veen JA, Lemos EGM, Kuramae EE. 2017. Characterization of novel Acidobacteria exopolysaccharides with potential industrial and ecological applications. Sci Rep 7:41193.

34. Kalam S, Basu A, Ahmad I, Sayyed RZ, El-Enshasy HA, Dailin DJ, Suriani NL. 2020. Recent Understanding of Soil Acidobacteria and Their Ecological Significance: A Critical Review. Frontiers in Microbiology 11:2712.

35. Trojan D, García-Robledo E, Hausmann B, Revsbech NP, Woebken D, Eichorst SA. 2024. A respiro-fermentative strategy to survive nanoxia in Acidobacterium capsulatum. FEMS Microbiol Ecol 100:fiae152.

36. Costa OYA, Zerillo MM, Zühlke D, Kielak AM, Pijl A, Riedel K, Kuramae EE. 2020. Responses of Acidobacteria Granulicella sp. WH15 to High Carbon Revealed by Integrated Omics Analyses. Microorganisms 8:244.

37. Halamka TA, Garcia A, Evans TW, Schubert S, Younkin A, Hinrichs K-U, Kopf S. 2024. Occurrence of ceramides in the Acidobacterium Solibacter usitatus: implications for bacterial physiology and sphingolipids in soils. Front Geochem 2.

38. Sinninghe Damsté JS, Rijpstra WIC, Hopmans EC, Weijers JWH, Foesel BU, Overmann J, Dedysh SN. 2011. 13,16-Dimethyl Octacosanedioic Acid (iso-Diabolic Acid), a Common Membrane-Spanning Lipid of Acidobacteria Subdivisions 1 and 3. Applied and Environmental Microbiology 77:4147–4154.

39. Oberpaul M, Brinkmann S, Marner M, Mihajlovic S, Leis B, Patras MA, Hartwig C, Vilcinskas A, Hammann PE, Schäberle TF, Spohn M, Glaeser J. 2022. Combination of high-throughput microfluidics and FACS technologies to leverage the numbers game in natural product discovery. Microbial Biotechnology 15:415–430.

40. Chklovski A, Parks DH, Woodcroft BJ, Tyson GW. 2023. CheckM2: a rapid, scalable and accurate tool for assessing microbial genome quality using machine learning. Nat Methods 20:1203–1212.

41. Bowers RM, Kyrpides NC, Stepanauskas R, Harmon-Smith M, Doud D, Reddy TBK, Schulz F, Jarett J, Rivers AR, Eloe-Fadrosh EA, Tringe SG, Ivanova NN, Copeland A, Clum A, Becraft ED, Malmstrom RR, Birren B, Podar M, Bork P, Weinstock GM, Garrity GM, Dodsworth JA, Yooseph S, Sutton G, Glöckner FO, Gilbert JA, Nelson WC, Hallam SJ, Jungbluth SP, Ettema TJG, Tighe S, Konstantinidis KT, Liu W-T, Baker BJ, Rattei T, Eisen JA, Hedlund B, McMahon KD, Fierer N, Knight R, Finn R, Cochrane G, Karsch-Mizrachi I, Tyson GW, Rinke C, Lapidus A, Meyer F, Yilmaz P, Parks DH, Murat Eren A, Schriml L, Banfield JF, Hugenholtz P, Woyke T. 2017. Minimum information about a single amplified genome (MISAG) and a metagenome-assembled genome (MIMAG) of bacteria and archaea. Nat Biotechnol 35:725–731.

42. Parks DH, Chuvochina M, Rinke C, Mussig AJ, Chaumeil P-A, Hugenholtz P. 2022. GTDB: an ongoing census of bacterial and archaeal diversity through a phylogenetically consistent, rank normalized and complete genome-based taxonomy. Nucleic Acids Research 50:D785–D794.

43. Bode HB, Bethe B, Höfs R, Zeeck A. 2002. Big effects from small changes: possible ways to explore nature’s chemical diversity. Chembiochem:6 –627.

44. Sumner LW, Amberg A, Barrett D, Beale MH, Beger R, Daykin CA, Fan TW-M, Fiehn O, Goodacre R, Griffin JL, Hankemeier T, Hardy N, Harnly J, Higashi R, Kopka J, Lane AN, Lindon JC, Marriott P, Nicholls AW, Reily MD, Thaden JJ, Viant MR. 2007. Proposed minimum reporting standards for chemical analysis Chemical Analysis Working Group (CAWG) Metabolomics Standards Initiative (MSI). Metabolomics 3:211–221.

45. Letunic I, Bork P. 2019. Interactive Tree Of Life (iTOL) v4: recent updates and new developments. Nucleic Acids Res 47:W256–W259.

46. Blin K, Shaw S, Kloosterman AM, Charlop-Powers Z, van Wezel GP, Medema MH, Weber T. 2021. antiSMASH 6.0: improving cluster detection and comparison capabilities. Nucleic Acids Res 49:W29–W35.

47. Navarro-Muñoz JC, Selem-Mojica N, Mullowney MW, Kautsar SA, Tryon JH, Parkinson EI, De Los Santos ELC, Yeong M, Cruz-Morales P, Abubucker S, Roeters A, Lokhorst W, Fernandez-Guerra A, Cappelini LTD, Goering AW, Thomson RJ, Metcalf WW, Kelleher NL, Barona-Gomez F, Medema MH. 2020. A computational framework to explore large-scale biosynthetic diversity. Nat Chem Biol 16:60–68.

48. Gu G, Smith L, Liu A, Lu S-E. 2011. Genetic and Biochemical Map for the Biosynthesis of Occidiofungin, an Antifungal Produced by Burkholderia contaminans Strain MS14 ▿. Appl Environ Microbiol 77:6189–6198.

49. Cothrell A, Cao K, Bonasera R, Tenorio A, Orugunty R, Smith L. 2023. Intravaginal Gel for Sustained Delivery of Occidiofungin and Long-Lasting Antifungal Effects. Gels 9:787.

50. Spaepen S, Vanderleyden J, Remans R. 2007. Indole-3-acetic acid in microbial and microorganism-plant signaling. FEMS Microbiology Reviews 31:425–448.

51. Schmidt CS, Mrnka L, Frantík T, Lovecká P, Vosátka M. 2018. Plant growth promotion of iscanthus × giganteus by endophytic bacteria and fungi on non-polluted and polluted soils. World J Microbiol Biotechnol 34:48.

52. Lovecká P, Kroneislová G, Novotná Z, Röderová J, Demnerová K. 2023. Plant Growth-Promoting Endophytic Bacteria Isolated from Miscanthus giganteus and Their Antifungal Activity. Microorganisms 11:2710.

53. Santana MM, Rosa AP, Zamarreño AM, García-Mina JM, Rai A, Cruz C. 2022. Achromobacter xylosoxidans and Enteromorpha intestinalis Extract Improve Tomato Growth under Salt Stress. Agronomy 12:934.

54. Environment Canada, Health Canada. 2015. Final Screening Assessment for Paenibacillus polymyxa ATCC 842; Paenibacillus polymyxa ATCC 55407; Paenibacillus polymyxa 13540-4. En14-230/2015E-PDF. Final Screening Assessment. Government of Canada, Ottawa.

55. Yoneda Y, Yamamoto K, Makino A, Tanaka Y, Meng X-Y, Hashimoto J, Shin-ya K, Satoh N, Fujie M, Toyama T, Mori K, Ike M, Morikawa M, Kamagata Y, Tamaki H. 2021. Novel Plant-Associated Acidobacteria Promotes Growth of Common Floating Aquatic Plants, Duckweeds. Microorganisms 9:1133.

56. Akhtar SS, Mekureyaw MF, Pandey C, Roitsch T. 2020. Role of Cytokinins for Interactions of Plants With Microbial Pathogens and Pest Insects. Front Plant Sci 10.

57. Fu S-F, Wei J-Y, Chen H-W, Liu Y-Y, Lu H-Y, Chou J-Y. 2015. Indole-3-acetic acid: A widespread physiological code in interactions of fungi with other organisms. Plant Signal Behav 10:e1048052.

58. Koenig RL, Morris RO, Polacco JC. 2002. tRNA Is the Source of Low-Level trans-Zeatin Production in Methylobacterium spp. Journal of Bacteriology 184:1832–1842.

59. Wei X, Moreno-Hagelsieb G, Glick BR, Doxey AC. 2023. Comparative analysis of adenylate isopentenyl transferase genes in plant growth-promoting bacteria and plant pathogenic bacteria. Heliyon 9:e13955.

60. Frébortová J, Frébort I. 2021. Biochemical and Structural Aspects of Cytokinin Biosynthesis and Degradation in Bacteria. Microorganisms 9:1314.

61. Sugawara H, Ueda N, Kojima M, Makita N, Yamaya T, Sakakibara H. 2008. Structural insight into the reaction mechanism and evolution of cytokinin biosynthesis. Proc Natl Acad Sci U S A 105:2734–2739.

62. Kuroha T, Tokunaga H, Kojima M, Ueda N, Ishida T, Nagawa S, Fukuda H, Sugimoto K, Sakakibara H. 2009. Functional Analyses of LONELY GUY Cytokinin-Activating Enzymes Reveal the Importance of the Direct Activation Pathway in Arabidopsis. Plant Cell 21:3152–3169.

63. Seo H, Kim S, Sagong H-Y, Son HF, Jin KS, Kim I-K, Kim K-J. 2016. Structural basis for cytokinin production by LOG from Corynebacterium glutamicum. Sci Rep 6:31390.

64. Noor A, Ziaf K, Naveed M, Khan KS, Ghani MA, Ahmad I, Anwar R, Siddiqui MH, Shakeel A, Khan AI. 2023. L-Tryptophan-Dependent Auxin-Producing Plant-Growth-Promoting Bacteria Improve Seed Yield and Quality of Carrot by Altering the Umbel Order. Horticulturae 9:954.

65. Kravchenko LV, Azarova TS, Makarova NM, Tikhonovich IA. 2004. The Effect of Tryptophan Present in Plant Root Exudates on the Phytostimulating Activity of Rhizobacteria. Microbiology 73:156–158.

66. Eze MO, Amuji CF. 2024. Elucidating the significant roles of root exudates in organic pollutant biotransformation within the rhizosphere. Sci Rep 14:2359.

67. Fones H, Gurr S. 2015. The impact of Septoria tritici Blotch disease on wheat: An EU perspective. Fungal genetics and biology : F & B : – 7.

68. Peng J, Lin T, Wang W, Xin Z, Zhu T, Gu Q, Li D. 2013. Antiviral Alkaloids Produced by the Mangrove-Derived Fungus Cladosporium sp. PJX-41. J Nat Prod 76:1133–1140.

69. Mayser P, Wenzel M, Krämer H-J, Kindler BLJ, Spiteller P, Haase G. 2007. Production of indole pigments by Candida glabrata. Med Mycol 45:519–524.

70. Mayser P, Wille G, Imkampe A, Thoma W, Arnold N, Monsees T. 1998. Synthesis of fluorochromes and pigments in Malassezia furfur by use of tryptophan as the single nitrogen source. Mycoses 41:265–271.

71. Mayser P, Schäfer U, Krämer H-J, Irlinger B, Steglich W. 2002. Pityriacitrin – an ultraviolet-absorbing indole alkaloid from the yeast Malassezia furfur. Arch Dermatol Res 294:131–134.

72. Irlinger B, Bartsch A, Krämer H-J, Mayser P, Steglich W. 2005. New Tryptophan Metabolites from Cultures of the Lipophilic Yeast Malassezia furfur. Helvetica Chimica Acta 88:1472–1485.

73. Huang D, Zhang Z, Li Y, Liu F, Huang W, Min Y, Wang K, Yang J, Cao C, Gong Y, Ke S. 2022. Carboline derivatives based on natural pityriacitrin as potential antifungal agents. Phytochemistry Letters 48:100–105.

74. Machowinski A, Krämer H-J, Hort W, Mayser P. 2006. Pityriacitrin – a potent UV filter produced by Malassezia furfur and its effect on human skin microflora. Mycoses 49:388–392.

75. Dai J, Dan W, Schneider U, Wang J. 2018. β-Carboline alkaloid monomers and dimers: Occurrence, structural diversity, and biological activities. Eur J Med Chem 157:622–656.

76. Gaitanis G, Magiatis P, Mexia N, Melliou E, Efstratiou MA, Bassukas ID, Velegraki A. 2019. Antifungal activity of selected Malassezia indolic compounds detected in culture. Mycoses 62:597–603.

77. Wang J, Chen M, Lv Y, Jiang Y, Qiu L. 2016. Edaphobacter dinghuensis sp. nov., an acidobacterium isolated from lower subtropical forest soil. International Journal of Systematic and Evolutionary Microbiology, 66:276–282.

78. Ueda S, Kitani S, Namba T, Arai M, Ikeda H, Nihira T. 2018. Engineered production of kitasetalic acid, a new tetrahydro-β-carboline with the ability to suppress glucose-regulated protein synthesis. J Antibiot 71:854–861.

79. Li X-L, Sun Y, Yin Y, Zhan S, Wang C. 2023. A bacterial-like Pictet–Spenglerase drives the evolution of fungi to produce β-carboline glycosides together with separate genes. Proceedings of the National Academy of Sciences 120:e2303327120.

80. Zuther K, Mayser P, Hettwer U, Wu W, Spiteller P, Kindler BLJ, Karlovsky P, Basse CW, Schirawski J. 2008. The tryptophan aminotransferase Tam1 catalyses the single biosynthetic step for tryptophan-dependent pigment synthesis in Ustilago maydis. Molecular Microbiology 68:152–172.

81. Liew LPP, Fleming JM, Longeon A, Mouray E, Florent I, Bourguet-Kondracki M-L, Copp BR. 2014. Synthesis of 1-indolyl substituted β-carboline natural products and discovery of antimalarial and cytotoxic activities. Tetrahedron 70:4910–4920.

82. Khosravi Babadi Z, Ebrahimipour G, Wink J, Narmani A, Risdian C. 2021. Isolation and identification of Streptomyces sp. Act4Zk, a good producer of Staurosporine and some derivatives. Lett Appl Microbiol 72:206–218.

83. Patz S, Gautam A, Becker M, Ruppel S, Rodríguez-Palenzuela P, Huson D. 2021. PLaBAse: A comprehensive web resource for analyzing the plant growth-promoting potential of plant-associated bacteria. bioRxiv 10.1101/2021.12.13.472471.

84. Eichberg J, Oberpaul M, Hartwig C, Geißler AH, Culmsee C, Vilcinskas A, Böttcher-Friebertshäuser E, Brückner H, Degenkolb T, Hardes K. 2024. Structural characterization and bioactivity profiling of the fungal peptaibiotic tolypin reveal protective effects against influenza viruses. Archiv der Pharmazie 357:e2400384.

85. Brinkmann S, Kurz M, Patras MA, Hartwig C, Marner M, Leis B, Billion A, Kleiner Y, Bauer A, Toti L, Pöverlein C, Hammann PE, Vilcinskas A, Glaeser J, Spohn M, Schäberle TF. 2022. Genomic and Chemical Decryption of the Bacteroidetes Phylum for Its Potential to Biosynthesize Natural Products. Microbiology Spectrum 10:e02479–21.

