## Supplementary material for "Tryptophan-Driven Metabolomic Shift in *Acidobacteriaceae* Reveals Phytohormones and Antifungal Metabolites": SI_ExploringAcidobacteriota

### Extended Methods and Results

#### Methanolic Extract Generation

Lyophilised microbial cultures in 24-well plates were extracted with 3 ml methanol (MeOH) per well. Extracts were sonicated for five minutes, shaken (28°C, two h) and centrifuged (4000 rpm, room temperature (RT), 10 minutes). The supernatants were transferred to 96-well plates and dried completely by speed-vacuuming. Residues were dissolved in 200 µl MeOH, homogenised using a TissueLyzer II (Qiagen, Hilden, Germany) and by sonication as previously described. Plates were then stored overnight at 4°C for precipitation, followed by centrifugation. From this plate, two 96-well V-bottom plates were prepared: the Assay Master Plate, for screening purposes (100 µl of extract), and an analytics plate (70 µl for metabolite profiling by UPLC-HRMS).

Lyophilised microbial cultures in Erlenmeyer flasks were extracted with 40 mL MeOH by shaking at 230 rpm at 28°C for 2 hours. Extracts were transferred to Falcon tubes, centrifuged (4000 rpm, 15 min, RT), and filtered (30 µm). The supernatants were evaporated to dryness overnight. Dried extracts were redissolved in 1 ml MeOH, transferred to 1.5 ml Eppendorf tubes, incubated at 4°C overnight for precipitation, before being centrifuged (13 000 rpm, 30 min, RT) to obtain clear extracts. These extracts were transferred to 96-deep-well plates and centrifuged again (4000 rpm, 10 min, RT). From this plate, we prepared three different plates, one for metabolite profiling by UPLC-HRMS (60 µl), screening (150 µl), and for storage (600 µl).

#### Plant assays

Barley seedlings (Organic Grains "Summer Barley" (EAN 7640126481333) from Sativa Rheinau purchased at Samenhaus Müller GmbH) were first disinfected and treated with antibiotics to prevent the growth of other competing microorganisms. To do this, commercially available bleach (DanKlorix) was diluted with water to a final concentration of 25%. The required number of barley seeds was selected and transferred into a beaker. Seeds were covered with bleach solution and incubated for 15 minutes with continuous stirring. Afterwards, the seeds were rinsed 5 times with distilled water. Seeds that were floating in the water were removed from the sample. Finally, seeds were dried under a sterile bench on paper towels for five hours before being stored in a sterile 50 ml Falcon tube. For plant testing, sterile glass vials (Fisher Brand, Cat. Nr. 14-961-34) were filled with 5 mL of SH medium (Schenk and Hildebrandt Basal Salt Mixture, 3.2 g/l) and (pH5.8) containing 0.25% Gelzan (Gelrite) for solidification. The medium was then further supplemented with Pen/Strep Solution (final concentration: 1 U/µl penicillin and 1 µg/ml Streptomycin) and Nystatin (final concentration: 0.64 µg/ml) to prevent bacterial and fungal growth. After the liquid had evaporated, the barley seeds were transferred to the medium and covered with a sterile plastic cover (Kuble Art. No. 73660-25, 25 mm). Vials with seeds were incubated in the growth chamber for 72 hours in the dark before switching to a day and night cycle with intervals of 16 hours light and 8 hours dark at 24°C and 18°C, respectively. We grew seeds for 6 days before the final evaluation with random positioning every day. For the final assessment, the vials were boiled to liquefy the medium, the medium was washed off the roots, the roots were separated from the seedlings, and the seed and lyophilised to determine the dry weight of the different plant components.

#### Analysis of assemblies and their biosynthetic gene cluster potential

The 618 curated genome assemblies were processed to comparatively evaluate metadata such as assembly size, GC-content and N50, monitored by Geneious (v11.1.5), and to predict the BGC potential. BGC prediction was performed using antiSMASH (v6.1.1); afterwards, predicted BGCs were analysed using BiG-SCAPE (1) at a cutoff of 0.5, including MiBIG reference clusters in the analysis. The sequence similarity network was visualised using Cytoscape (v3.10.2). UMAP was used to reduce the high-dimensional biosynthetic similarity data reported by BiG-SCAPE into two dimensions, allowing for the visualisation of overall differences in BGC composition between different strains in our acidobacterial dataset. The binary presence/absence matrix (BiG-SCAPE output) was used to compute pairwise cosine similarities using scikit-learn (version 1.3). The matrix of cosine distances was then projected in two dimensions using UMAP (UMAP-learn, v0.5.5, metric='cosine', random_state=42) and visualised using matplotlib (v3.7). Strains were coloured according to their taxonomic class to reflect phylogenetic relationships.

Our analysis revealed that the metagenome-assembled genomes (MAGs) are, on average, slightly smaller (4.88 ± 1.46 Mb) than isolate genomes (5.28 ± 1.71 Mb) (Figure S4). This difference is statistically modest with the Mann–Whitney U test yielding $p = 0.04$, which is significant at the 5% level but does not meet a more stringent 1% threshold. All assemblies not affiliated to a genus by GTDB are MAGs (n=85; 13.7%). Contig numbers ranged from 1 to 1357, with an average of 200 contigs per genome. The BGC number did not significantly correlate with the contig number, indicating no significant overestimation of BGCs due to fragmented clusters (Figure S7). Interestingly, the BGC number also did not correlate with the assembly size, a phenomenon typically observed across different taxonomic levels (2–4).

The GC content of the analysed genome set ranged from 34.3% to 73.8% (Figure S5). This is substantially broader than previously proposed for the Acidobacteriota phylum (5). This broad GC range exceeds the typical GC distribution reported for most bacterial phyla, with comparable broadness covered only by the *Pseudomonadota*. Notably, from our dataset, 37 genomes exhibited a GC content ≥70%, all belonging to the three classes Mor1 (average GC% 68.0 ± 4.2, n9), *Vicinamibacteria* (average GC% 67.5 ± 2.7, n98), and *Thermoanaerobaculia* (average GC% 67.9 ± 3.2, n79) (Figure S6). GC-rich bacterial genomes are considered an adaptation to high temperatures, with a demonstrated positive correlation between the optimal growth temperature of bacteria (6–8) and their GC content. Indeed, the *Thermoanaerobaculia* class has been generally associated with high-temperature environments, and its single isolated strain, *Thermoanaerobaculum aquaticum* DSM 24856, exhibits optimal growth at 60°C (9). Considering the wide range of GC-content of the Acidobacteriota phylum, we evaluated whether the GC-content correlates with the number of BGCs detected, but did not find any correlation (Figure S5; 0.03; $p = 0.482$). Indeed, the most enriched classes in terms of GC content (Mor1, *Vicinamibacteria*, *Thermoanaerobaculia*; Figure S6) are rather under-equipped in BGC load, and only Mor1 shows a moderate correlation between GC content and BGC number. Instead, the *Acidobacteriae* class exhibits a strong negative correlation, with lower GC-content genomes harbouring a larger number of BGCs. The most recognised bacterial NP producing taxa, the Actinomycetes class of the Actinomycetota phylum, is well known for its large genomes (up to 10 Mbps) and high GC content (10). Within this phylum, genome size and GC content appear to be phylogenetically conserved and positively correlated (11). Furthermore, genome size and number of secondary metabolite biosynthetic gene clusters are positively correlated in the Actinomycetes class, indicating that larger genomes can accommodate more gene clusters devoted to secondary metabolism

(12). The most talented orders for natural product biosynthesis (e.g. *Streptomycetales*, *Pseudonocardiales*, *Streptosporangiales* and *Micromonosporales*) carry the biggest genomes (average > 7.5 kb) and highest GC content (average >70%), outcompeting orders with lower biosynthetic potential (e.g. *Corynebacteriales*, *Propionibacteriales*, *Micrococcales*, and *Bifidobacteriales*) characterised by comparatively low genome sizes (average < 5.1 kb) and lower GC content (average < 70%) (11).

Evaluation of the Gene Cluster Families (GCFs) generated by BiG-SCAPE (1), enabled conclusions on their distributions within the dataset and their correlation to BGCs, encoding the production of known NPs. Filtering revealed GCFs shared among different acidobacterial classes (n = 22; Figure S10). Most of the GCFs are shared between the *Acidobacteriae* and the *Vicinamibacteria* (n = 7). There is no clear trend in BGC cluster type; instead, NRP, RiPP, Terpenes, and PKS1 cluster types are shared. In general, the class of *Acidobacteriae* shares GCFs with every class except for UBA890 and Mor1. The most diverse GCFs when it comes to classes are GCF3545 and GCF5459, both RiPP clusters shared between three different classes (*Vicinamibacteria* and *Acidobacteriae* and *Blastocatellia* or UBA6911, respectively). The last one is GCF5622, a PKS1 cluster harboured by *Acidobacteriae*, UBA6911 and *Vicinamibacteria*.

GCF alignment to the MiBIG Reference BGCs revealed several matches. In total, 11 acidobacterial clusters, grouped into six different GCFs, showed similarities to known MiBIG reference clusters. However, CORASON alignment of these clusters identified two GCFs initially matched to MiBIG reference clusters that result from fragmented PKS and NRPS genes, located at contig edges, preventing complete cluster reconstruction. These represent a PKS1-like fragment associated with aromatic/benzenoid structures (GCF1211) and an NRPS fragment with partial similarity to anabaenopeptin-type clusters (GCF4479) (not shown in Figure 2). The remaining four confirmed clusters span various compound classes and taxonomic origins. GCF1752 was associated with alkylpyrone-407 and alkylpyrone-393, and a similar cluster, present in the *Thermoanaerobaculia* class, suggesting the production of small, polyketide-derived alkylated heterocycles. Besides PKS systems, we identified three NRP GCFs with related, known MiBIG reference clusters. GCF1359 contains different cluster types, all of which produce myxochromide D and S, linked to an NRP cluster from a representative of the *Chloracidobacteriales*. One additional correlation hit was found within GCF1959, covering a PKS-NRPS hybrid of *Acidobacterium capsulatum* and the soil isolate FHG110511, which clusters with the occidiofungin A-reference BGC (MiBIG ID: BGC0001711) (13). We also found three input genomes, belonging to the classes *Vicinamibacteria* and *Acidobacteriae*, that clustered with the reference clusters for ectoine. The two incompletely

#### Genomic Analysis of PGPT Traits

DIAMOND was run as follows:

```
> diamond blastp -q <input.fasta> -d <PGPT_DB> -o <output.tsv>

--outfmt 6 qtitle qseqid sseqid pident length mismatch gapopen qstart qend sstart send evalue bitscore

--evalue 1e-5 --threads 8 --<sensitivity_mode>
```

DIAMOND results were further processed to include relevant metadata and to minimise redundant hits. First (1.) we retained hits with E-values ≤1E-50 (RawHits). Second, we filtered for the best hit per plant-growth-promoting-trait (PGPT) family for each CDS. Finally, we reduced the dataset to contain only the best overall PGPT hit per CDS based on the lowest e-value (Output labelled with prefix “best_”). This output was annotated by merging the filtered hits with the PGPT ontology file, thereby matching the PGPT family numbers with their hierarchical classifications and functions. The output was used to analyse gene-specific content of individual strains. An overall summary of PGPT families per analysed genome was logged in PGPT-summary.txt. To get insights into taxonomic distributions of PGPTs, we merged the results with taxonomic metadata previously generated from GTDB-Tk. Based on taxonomic affiliation, the PGPT output can be optionally grouped by specifying a rank (e.g., family, order).

To evaluate the PGPT potential at the genome level, we computed the functional coverage across the different PGPT hierarchy levels. For each genome or taxonomic batch, we considered the number of PGPT families detected, the total possible number of PGPT families present in the database per hierarchy level, to calculate:

$$Coverage = \left(\frac{Detected\ PGPTs_i}{Total\ Possible\ PGPTs}\right) \times 100$$

Where i is the genome or taxonomy group.

For taxonomic batch comparison, the individually calculated coverage per genome is averaged across all genomes within the batch.

$$Coverage\ = \frac{1}{N}\sum_{i=1}^{N}\left(\frac{Detected\ PGPTs_i}{Total\ PGPTs} \times 100\right)$$

The values were exported to an Excel file, where the different sheets cover different PGPT levels.

All analysis steps were performed in Python 3.9 using pandas, matplotlib, seaborn, and BioPython. DIAMOND v2 was used for protein alignment.

#### Analytical procedures

The generated high-resolution mass spectrometry (HR-MS) data sets were copied and processed with both line spectra thresholds (5,000 and 10,000). Bucketing was performed on both thresholds at the same time, resulting in one table representing the OSMAC process and a second table representing the Trp-induction experiments, with both tables containing all buckets deemed identical for every sample (under both thresholds). These tables were subsequently curated: Buckets were only deemed present if they were detected in the 10,000 samples (4,406 (OSMAC) / 3,673 buckets (Trp-Induction) did not meet this criterion and were deleted). To avoid "uniqueness" due to compounds being just above the detection threshold, entries from the 5,000 set were used for these buckets; the 10,000 entries were deleted. 37 (OSMAC) / 54 (Trp-Induction) buckets were not filled in the respective tables and were subsequently deleted, resulting in two final tables representing 219 samples with 3,092 buckets (OSMAC) and 48 samples with 2,759 buckets (Trp-Induction), respectively. For OSMAC-barcoding, the conditions of each strain-media combination were combined. Each feature was deemed present when it occurred in two (grey) or more (black) extracts of the strain-media combination. Features that occurred only once were ignored as noise. Media control extracts were produced twice for each chemical composition (except for two media, where n = 1). For each chemical composition, a feature was deemed present when it occurred once (grey) or twice (black). The final table contained 2747 buckets, which were used for barcode generation.

Instead, for the Trp-Induction, a logical analysis was performed. Buckets filled in samples without tryptophan were coloured grey if present. For samples spiked with tryptophan, the sample was compared to its non-spiked counterpart and each bucket was categorised and assigned as Extinguished; Strong increase; Strong reduction; Light reduction; Present; Light increase; Strong increase:

| Non-Spiked | Spiked | Ratio Spiked/Non-Spiked | Category |
|---|---|---|---|
| Present | Absent | | Extinguished |
| Absent | Present | | Strong increase |
| Present | Present | > 0 and < 0.5 | Strong reduction |
| Present | Present | ≥ 0.5 and < 0.8 | Light reduction |
| Present | Present | ≥ 0.8 and < 1.25 | Present |
| Present | Present | ≥ 1.25 and < 2.0 | Light increase |
| Present | Present | ≥ 2.0 | Strong increase |

To achieve a readable representation, the buckets in the input table were sorted "decreasing" according to presence/absence for each sample, then according to "sum present" and subsequently for "present in media". Readers need to be aware that the colouration of buckets categorised as "extinguished" does not mean they are present anymore in the Trp-spiked sample.

### References


1. Navarro-Muñoz JC, Selem-Mojica N, Mullowney MW, Kautsar SA, Tryon JH, Parkinson EI, De Los Santos ELC, Yeong M, Cruz-Morales P, Abubucker S, Roeters A, Lokhorst W, Fernandez-Guerra A, Cappelini LTD, Goering AW, Thomson RJ, Metcalf WW, Kelleher NL, Barona-Gomez F, Medema MH. 2020. A computational framework to explore large-scale biosynthetic diversity. Nat Chem Biol 16:60–68.

2. Brinkmann S, Kurz M, Patras MA, Hartwig C, Marner M, Leis B, Billion A, Kleiner Y, Bauer A, Toti L, Pöverlein C, Hammann PE, Vilcinskas A, Glaeser J, Spohn M, Schäberle TF. 2022. Genomic and Chemical Decryption of the Bacteroidetes Phylum for Its Potential to Biosynthesize Natural Products. Microbiology Spectrum 10:e02479-21.

3. Männle D, McKinnie SMK, Mantri SS, Steinke K, Lu Z, Moore BS, Ziemert N, Kaysser L. 2020. Comparative Genomics and Metabolomics in the Genus Nocardia. mSystems 5:10.1128/msystems.00125-20.

4. Adamek M, Alanjary M, Sales-Ortells H, Goodfellow M, Bull AT, Winkler A, Wibberg D, Kalinowski J, Ziemert N. 2018. Comparative genomics reveals phylogenetic distribution patterns of secondary metabolites in Amycolatopsis species. BMC Genomics 19:426.

5. McReynolds E, Elshahed MS, Youssef NH. 2025. An ecological-evolutionary perspective on the genomic diversity and habitat preferences of the Acidobacteriota. Microbial Genomics 11:001344.

6. Hu E-Z, Lan X-R, Liu Z-L, Gao J, Niu D-K. 2022. A positive correlation between GC content and growth temperature in prokaryotes. BMC Genomics 23:110.

7. Lightfield J, Fram NR, Ely B. 2011. Across Bacterial Phyla, Distantly-Related Genomes with Similar Genomic GC Content Have Similar Patterns of Amino Acid Usage. PLOS ONE 6:e17677.

8. Teng W, Liao B, Chen M, Shu W. 2022. Genomic Legacies of Ancient Adaptation Illuminate GC-Content Evolution in Bacteria. Microbiology Spectrum 11:e02145-22.

9. Losey NA, Stevenson BS, Busse H-J, Damsté JSS, Rijpstra WIC, Rudd S, Lawson PA. 2013. Thermoanaerobaculum aquaticum gen. nov., sp. nov., the first cultivated member of Acidobacteria subdivision 23, isolated from a hot spring. International Journal of Systematic and Evolutionary Microbiology 63:4149–4157.

10. Barka EA, Vatsa P, Sanchez L, Gaveau-Vaillant N, Jacquard C, Klenk H-P, Clément C, Ouhdouch Y, van Wezel GP. 2015. Taxonomy, Physiology, and Natural Products of Actinobacteria. Microbiology and Molecular Biology Reviews 80:1–43.

11. Nouioui I, Carro L, García-López M, Meier-Kolthoff JP, Woyke T, Kyrpides NC, Pukall R, Klenk H-P, Goodfellow M, Göker M. 2018. Genome-Based Taxonomic Classification of the Phylum Actinobacteria. Front Microbiol 9.

12. Doroghazi JR, Metcalf WW. 2013. Comparative genomics of actinomycetes with a focus on natural product biosynthetic genes. BMC Genomics 14:611.

13. Terlouw BR, Blin K, Navarro-Muñoz JC, Avalon NE, Chevrette MG, Egbert S, Lee S, Meijer D, Recchia MJJ, Reitz ZL, van Santen JA, Selem-Mojica N, Tørring T, Zaroubi L, Alanjary M, Aleti G, Aguilar C, Al-Salihi SAA, Augustijn HE, Avelar-Rivas JA, Avitia-Domínguez LA, Barona-Gómez F, Bernaldo-Agüero J, Bielinski VA, Biermann F, Booth TJ, Carrion Bravo VJ, Castelo-Branco R, Chagas FO, Cruz-Morales P, Du C, Duncan KR, Gavriilidou A, Gayrard D, Gutiérrez-García K, Haslinger K, Helfrich EJN, van der Hooft JJJ, Jati AP, Kalkreuter E, Kalyvas N, Kang KB, Kautsar S, Kim W, Kunjapur AM, Li Y-X, Lin G-M, Loureiro C, Louwen JJR, Louwen NLL, Lund G, Parra J, Philmus B, Pourmohsenin B, Pronk LJU, Rego A, Rex DAB, Robinson S, Rosas-Becerra LR, Roxborough ET, Schorn MA, Scobie DJ, Singh KS, Sokolova N, Tang X, Udwary D, Vigneshwari A, Vind K, Vromans SPJM, Waschulin V, Williams SE, Winter JM, Witte TE, Xie H, Yang D, Yu J, Zdouc M, Zhong Z, Collemare J, Linington RG, Weber T, Medema MH. 2023. MiBIG 3.0: a community-driven effort to annotate experimentally validated biosynthetic gene clusters. Nucleic Acids Research 51:D603–D610.

14. Liew LPP, Fleming JM, Longeon A, Mouray E, Florent I, Bourguet-Kondracki M-L, Copp BR. 2014. Synthesis of 1-indolyl substituted β-carboline natural products and discovery of antimalarial and cytotoxic activities. Tetrahedron 70:4910–4920.

### Tables

Table S1 Bacterial strains and sources

| Strains | Source | Genome Information |
|---|---|---|
| *Edaphobacter aggregans* DSM 19364 | DSMZ | GCF_000745965.1; High contamination value – not in data set |
| *Acidobacterium sp.* S8 | From Petr Baldarian | GCF_009765985.1; High contamination value – not in dataset |
| *Silvibacterium bohemicum* S15 | From Petr Baldarian | GCF_001006305.1, quality fine, maybe should be included in the set |
| *Acidicapsa borealis* DSM 23886 | DSMZ | In-house genome |
| *Granulicella rosea* DSM 18704 | DSMZ | GCF_900188085.1, low completeness level |
| *Bryocella elongata* DSM 22489 | From Svetlana Dedysh | GCF_900108185.1, good quality genome, included in analysis |
| *FhG110202: Acidobacterium* | In-house strain, termite nest | In-house genome |
| *FhG110214:Terracidiphilus* | In-house strain, termite nest | In-house genome |
| *FhG110511:Edaphobacter* | In-house strain, soil | In-house genome |

Table S2 OSMAC media combinations

| Letter | Cultivation Size | Media composition | Media addition 1 | Media addition 2 |
|---|---|---|---|---|
| **A** | Small Scale | R2A pH 5.5 | | |
| **B** | Small Scale | 2x R2A pH 5.5 | | |
| **C** | Small Scale | 2x R2A pH 5.5 | | 0.05% GlcNAc |
| **D** | Small Scale | 2x R2A pH 5.5 | | 0.05% Pektin |
| **E** | Small Scale | 2x R2A pH 5.5 | | 0.5% Cellobiose |
| **F** | Small Scale | 2x R2A pH 5.5 | | 0.5% Xylan |
| **G** | Small Scale | 2x R2A pH 5.5 | | 0.8% Phytagel |
| **H** | Small Scale | 2x R2A pH 5.5 | 10 pcs Quartz beads | |
| **I** | Small Scale | 2x R2A pH 5.5 | | 3% Sucrose |
| **J** | Small Scale | 4x R2A pH 5.5 | | |
| **K** | Small Scale | HD pH 5.0 | | |
| **L** | Small Scale | HD pH 5.0 | 10 pcs Quartz beads | |
| **M** | Small Scale | HD pH 5.0 | 3 pcs ceramic beads | |
| **N** | Small Scale | 5294 pH 5.5 | | |

| | | | | |
|---|---|---|---|---|
| **O** | Small Scale | 5294 pH 5.5 | 10 pcs Quartz beads | |
| **P** | Small Scale | 5294 pH 5.5 | 3 pcs ceramic beads | |
| **Q** | Small Scale | PSYA5 pH 5.0 | | 0.5% Sucrose |
| **R** | Small Scale | PSYA5 pH 5.0 | 10 pcs Quartz beads | 0.5% Sucrose |
| **S** | Small Scale | PSYA5 pH 5.0 | 3 pcs ceramic beads | 0.5% Sucrose |
| **T** | Small Scale | PSYA5 pH 5.0 | | 3% Sucrose |
| **U** | Large Scale | VL55 | | 0.5% Xylane |
| **V** | Large Scale | VL55 | | 0.5% Sucrose |
| **W** | Large Scale | HD | | |
| **X** | Large Scale | 5294 modified | | |

**Table 3**: Summary of NMR analysis of Compound **1**

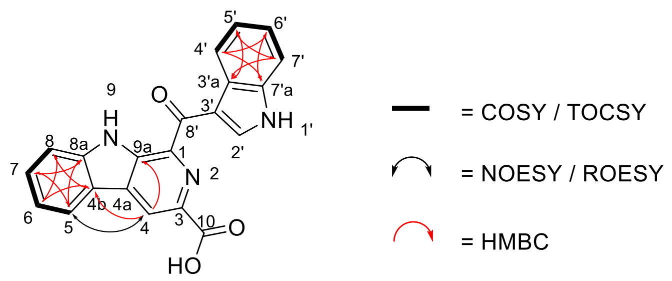


| Position | $\delta_C$ [ppm], Type | $\delta_H$ [ppm], mult. (J in Hz), int. |
|---|---|---|
| *1* | not observed | / |
| *2* | N | / |
| *3* | not observed | / |
| *4* | 118.9, CH[a] | 9.09, s |
| *4a* | not observed | / |
| *4b* | 120.4, $C_q$[b] | / |
| *5* | 122.1, CH[b] | 8.42, d (7.8) |
| *6* | 120.2, CH[a] | 7.34, t[c] (7.5) |
| *7* | 129.0, CH[b] | 7.64-7.60 (7.62)[d], m[e] |
| *8* | 113.3, CH[b] | 7.87, d (8.3), 1H |
| *8a* | 142.1, $C_q$[b] | / |
| *9* | NH | **12.36** / 12.25, br s |
| *9a* | 135.5, $C_q$[b] | / |
| *10* | not observed | / |
| *1'* | NH | 12.36 / **12.25**, br s |
| *2'* | not observed | not observed |
| *3'* | not observed | / |
| *3'a* | 127.5, $C_q$[b] | / |
| *4'* | 121.5, CH[b] | 8.61-8.56 (8.59)[d], m |
| *5'* | 122.1, CH[b] | 7.31-7.27 (7.29)[d], m |
| *6'* | 122.8, CH[b] | |
| *7'* | 112.2, CH[b] | 7.58-7.55 (7.56)[d], m |
| *7'a* | 136.3, $C_q$[b] | / |
| *8'* | not observed | / |

[a] The $^{13}$C Shift was extracted from the HSQC spectrum.

[b] The $^{13}$C Shifts was extracted from the HMBC spectrum.

[c] This peak might also be a non-resolved dd.

[d] $^{1}$H shift values in brackets denote the multiplet centers as extracted from the HSQC spectrum.

[e] The peak looks like a non-resolved dt.

**Table 4** Summary of NMR analysis of sample Compound 2

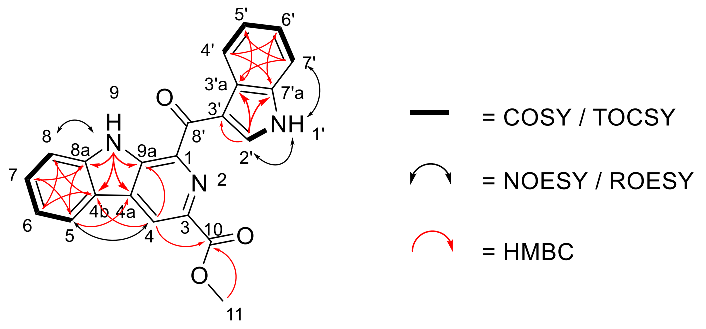


| *Position* | *δ<sub></sub>*$\delta_C$ *[ppm], Type* | $\delta_H$ *[ppm], mult. (J in Hz)* |
|---|---|---|
| *1* | 137.3, $C_q$[a] | / |
| *2* | N | / |
| *3* | 134.8, $C_q$[a] | / |
| *4* | 119.9, CH | 9.15, d (0.4) or s |
| *4a* | 131.3, $C_q$ | / |
| *4b* | 120.4, $C_q$ | / |
| *5* | 122.1, CH[b] | 8.48, d (7.9) |
| *6* | 120.8, CH | 7.38-7.36 (7.37)[c], m[d] |
| *7* | 129.2, CH | 7.65, ddd (8.2; 7.1; 1.1) |
| *8* | 113.4, CH | 7.90, d (8.2)[e] |
| *8a* | 142.1, $C_q$ | / |
| *9* | NH | 12.42, s |
| *9a* | 136.0, $C_q$ | / |
| *10* | 165.6, $C_q$ | / |
| *11* | 52.4, $CH_3$ | 4.04, s |
| *1’* | NH | 12.24, s |
| *2’* | 138.4, CH | 9.65, d (3.1) |
| *3’* | 114.1, $C_q$ | / |
| *3’a* | 127.2, $C_q$ | / |
| *4’* | 121.6, CH | 8.61-8.58 (8.59)[c], m |
| *5’* | 122.2, CH[b] | 7.32-7.28 (7.30)[c], m |
| *6’* | 123.0, CH | |
| *7’* | 112.3, CH | 7.59-7.57 (7.58)[c], m |
| *7’a* | 135.9, $C_q$ | / |
| *8’* | 186.3, $C_q$[a] | / |

[a] As no HMBC correlations were visible the signal was assigned by comparison with literature data (14).

[b] Distinction between C-5 and C-5’ according to literature.

[c] $^1$H shift values in brackets denote the multiplett centers as extracted from the HSQC spectrum.

[d] This peak is actually a ddd, but it is not resolved well enough to determine coupling constants.

[e] The peak looks like a non-resolved dt.

### Figures

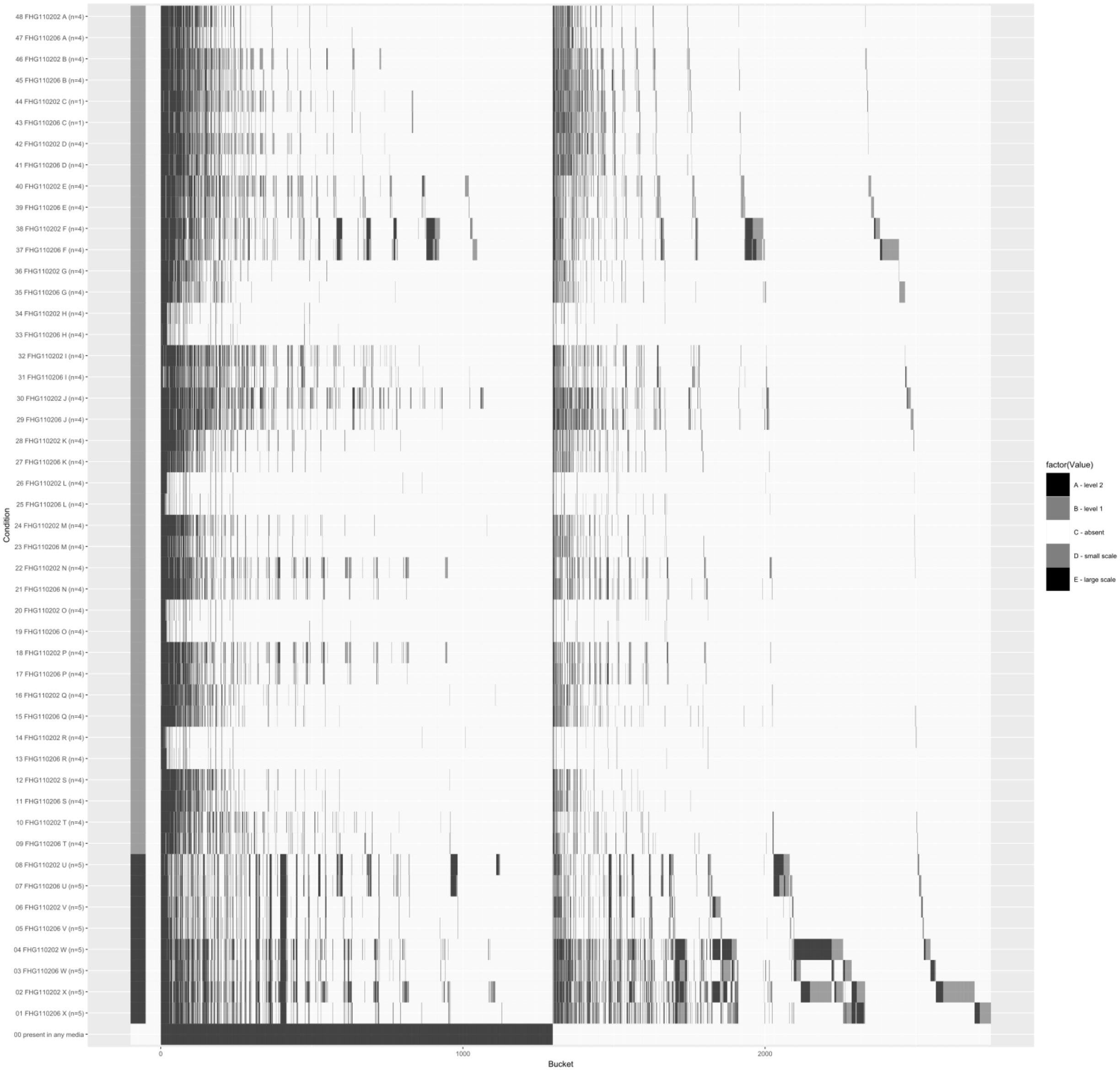


**Figure 1: UHPLC–QTOF–HR–MS profiles from 244 extracts of strains FHG110202 and FHG110206 (incl. media controls).** Profiles were aligned and bucketed into 2,747 features across 72 conditions. For strain–medium combinations, a bucket is considered present if it is observed in ≥2 extracts (grey = twice; black = ≥3); singletons were excluded. For media controls (columns, left, black label) (typically n=2), presence is scored once (grey) or twice (black). Quartz-beads (L, O, R, H) cultivations contained fewer buckets than the other conditions, while the large-scale cultivations (bottom, dark grey rows), showed more buckets compared to the small-scale cultivations (top, light grey rows).

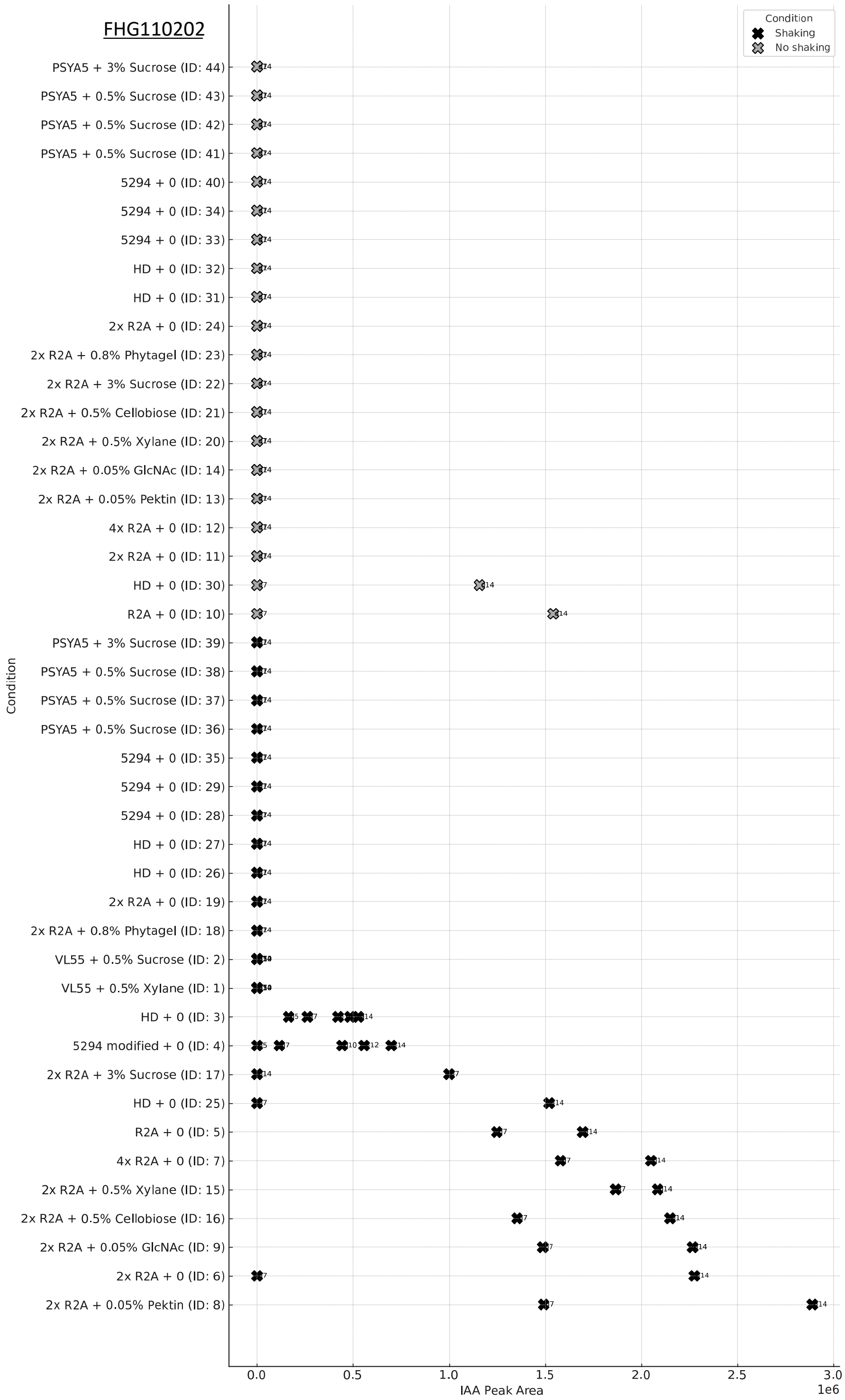


**Figure 2: Exemplary Production Profile of IAA (Raw Peak Area) in Strain FHG110202**. Feature 292.4s: 176.107m/z was extracted from the OSMAC feature table and tracked over time and condition (n =44, cf. OSMAC condition table for exact ID combinations, y-axis) to evaluate potential media composition and kinetics for further studies of IAA production (peak area, x-axis). Peak area values are plotted as crosses, with colour indicating agitation (light grey: No shaking; black: Shaking). Cultivation days are labelled beside crosses. In general, IAA production increases with cultivation time until day 14. The strain FHG110202 produced IAA in media base R2A, HD, and modified 5294, but not in PSYA5 and the original isolation medium VL55. Better production is observed with shaking rather than standing cultures and in R2A base medium.

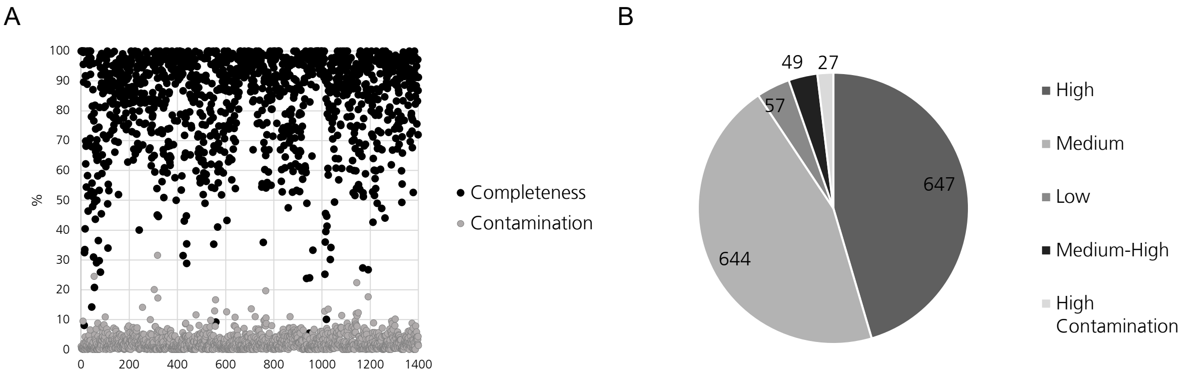


**Figure 3: CheckM2 Quality Control Data of NCBI-retrieved genomes labelled as belonging to the phylum Acidobacteriota.** A) Graph showing data from individual genomes indicating completeness (black) and contamination (grey) values in per cent. B) MIMAG evaluations of retrieved genomes. From 1428 genomes, we retained 647 high-quality genomes for further analysis.

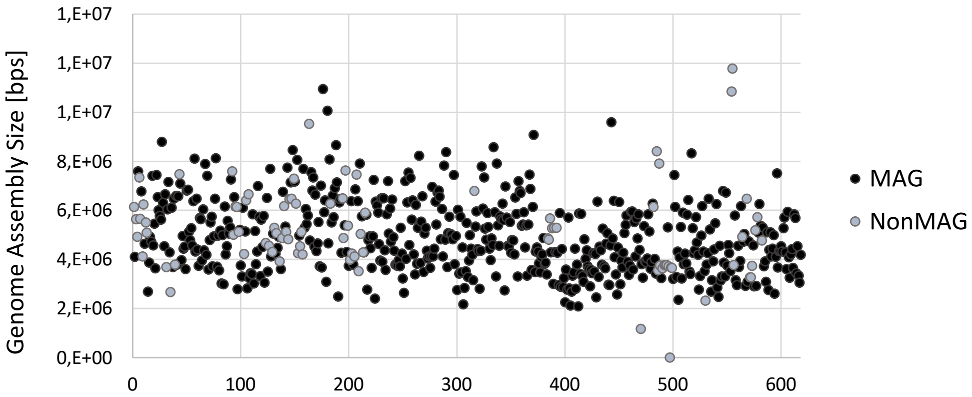


**Figure 4: Genome Assembly Size of Metagenome-assembled Genomes (black) and Isolate Genomes (Non-MAG, grey)**. Despite being slightly smaller on average, the genome size distribution of MAGs compared to that of isolate genomes is not statistically significant.

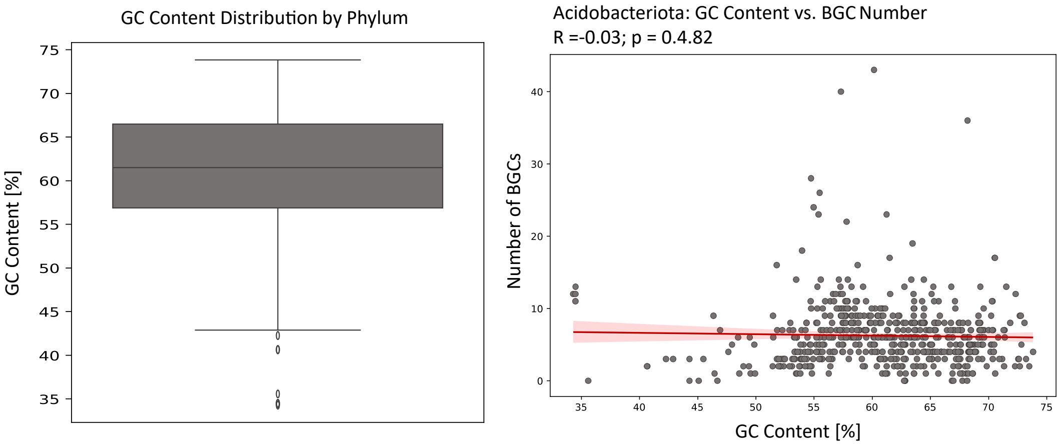


**Figure 5: Distribution of GC Content (left) and Correlation of GC Content and BGC Number (right) in the Acidobacteriota**.

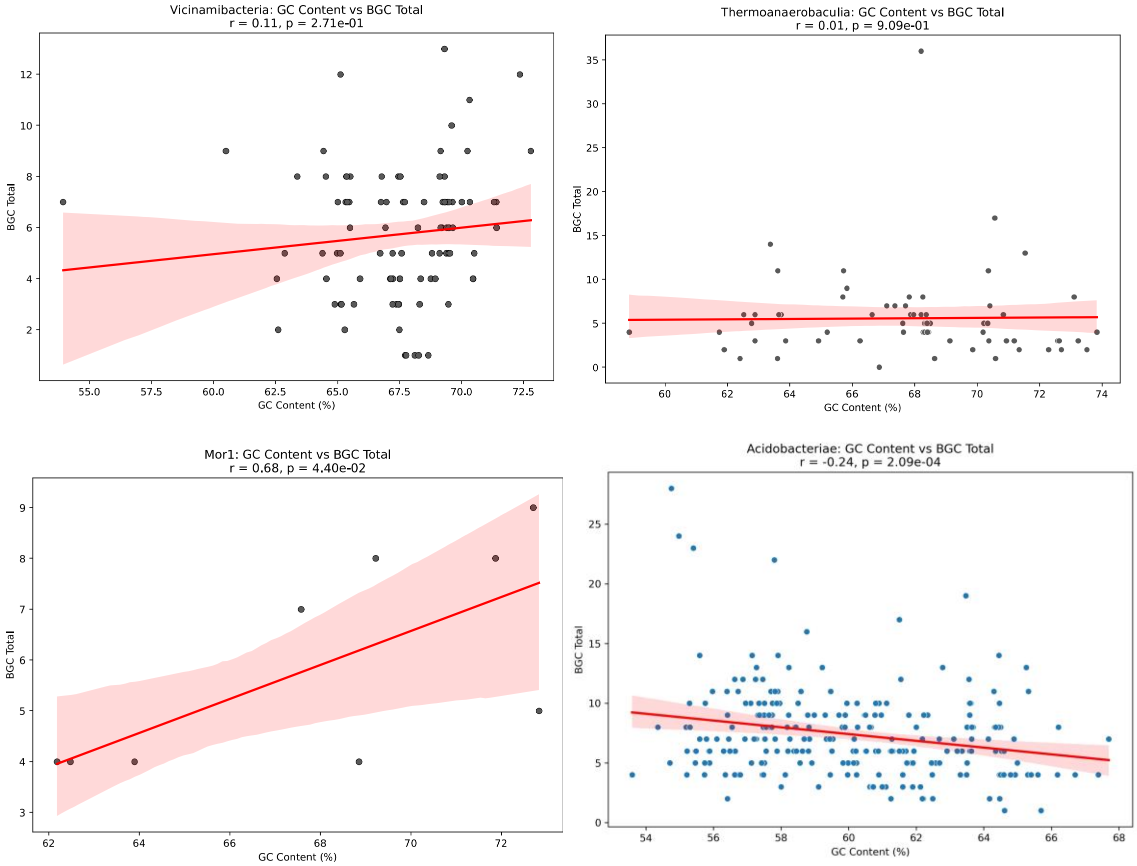


**Figure 6: Correlation of GC Content and BGC Number in acidobacterial classes (Mor1, *Vicinamibacteria*, *Thermoanaerobaculia* reaching 70% GC content and *Acidobacteriacea***. The Mor1 class shows higher BGC numbers with increasing GC content (p = 0.044), but exhibits an overall average BGC number compared to the rest of the phylum. The BGC numbers are higher in *Vicinamibacteria* and *Thermoanaerobaculia*, but there is no correlation between those and the GC content. In comparison, the class *Acidobacteriaceae* exhibits a clear negative correlation with genomes characterised by lower GC content, which harbour more BGCs.

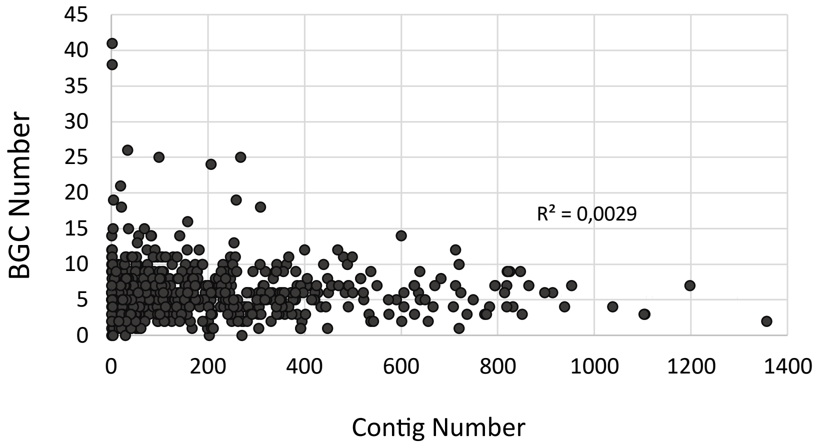


**Figure 7: BGC Number plotted against the Contig Number of Acidobacteriota Genomes.** There is no increase in BGC number with more fragmented genome assemblies (higher contig numbers), indicating that the total number of BGCs is not overestimated due to the presence of fragmented genome assemblies.

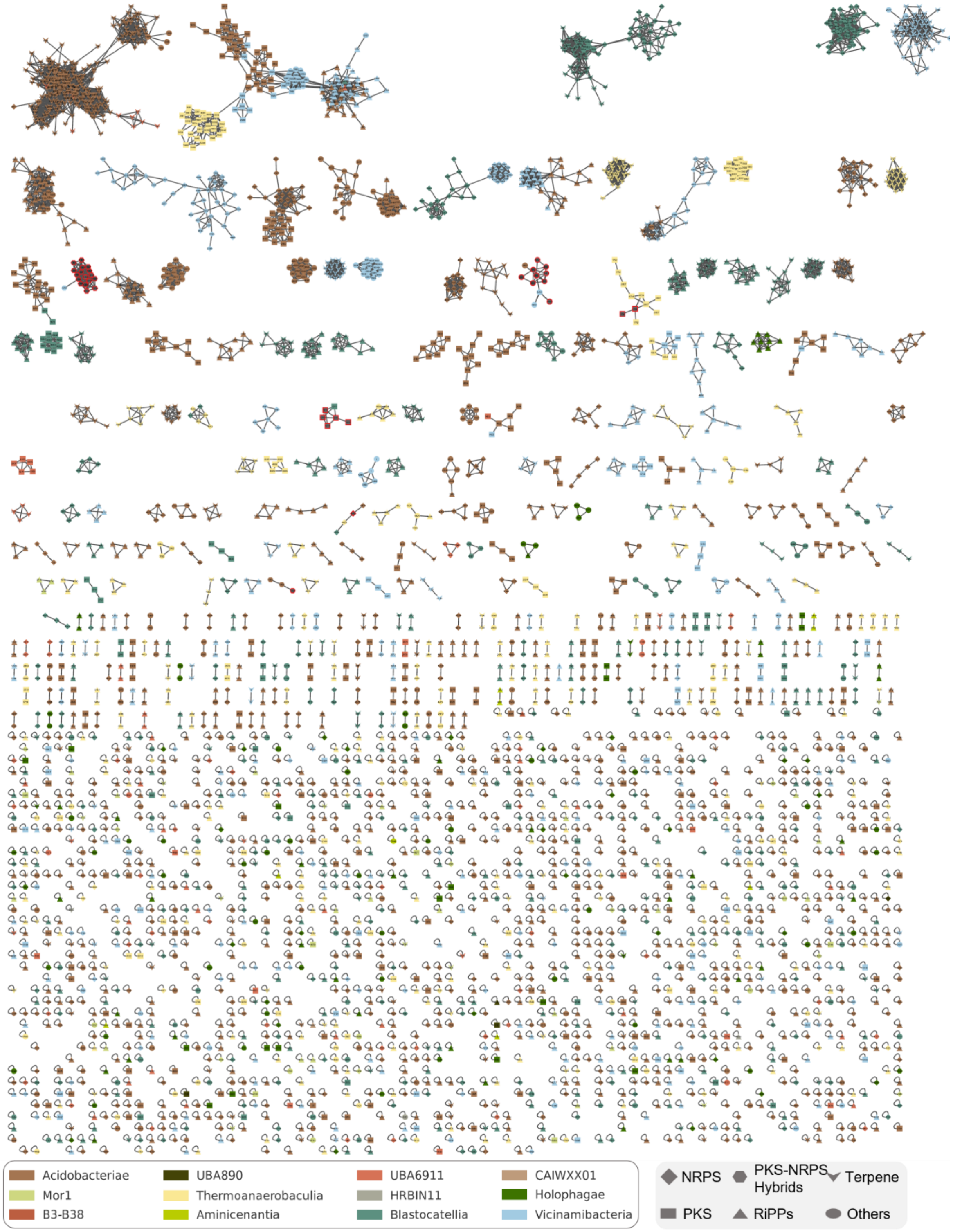


**Figure 8: BiG-SCAPE Similarity Network of antiSMASH-detected BGCs in the Acidobacteriota dataset at a cutoff value of 0.5.** Node colours indicate the taxonomic origin of the detected BGC at the class level (see legend, bottom left side). Node shapes indicate the BGC type assigned by (see legend, bottom right side). Nodes outlined in red correspond to MiBIG reference clusters.

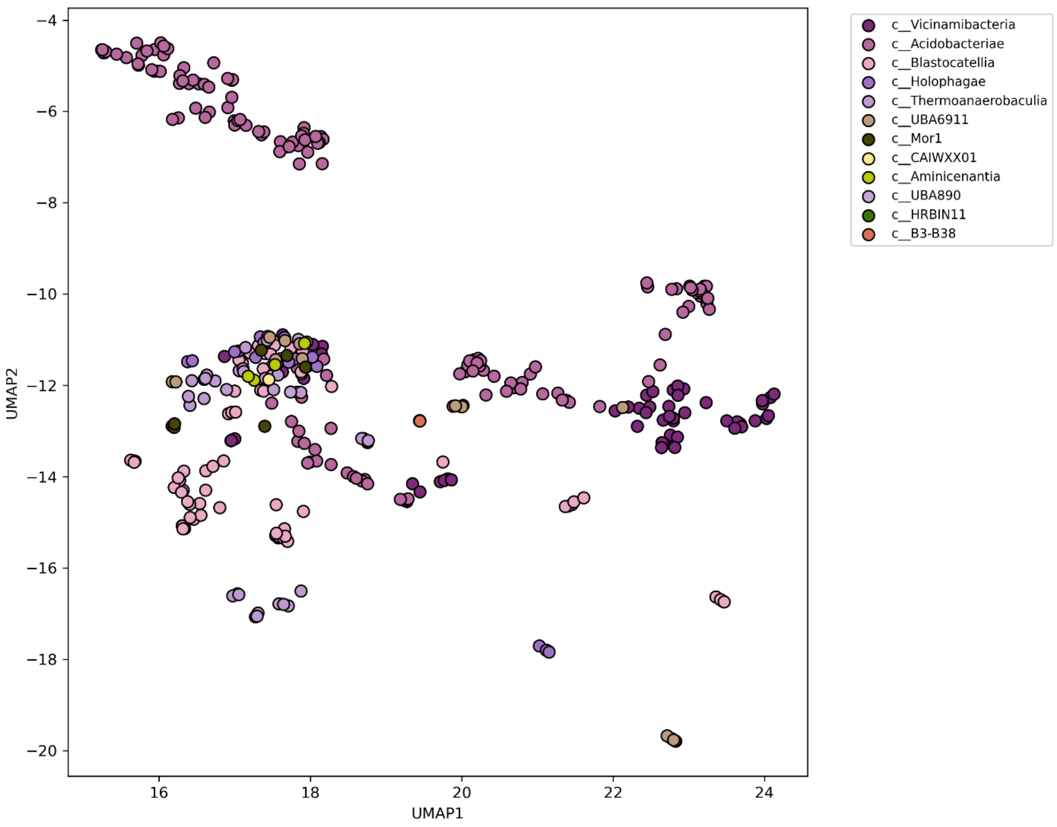


**Figure 9: UMAP Projection of Biosynthetic Gene Cluster (BGC) Profiles across Acidobacteriota Strains.** Each point represents a single microbial strain, positioned based on the cosine similarity of its biosynthetic gene cluster (BGC) composition using Uniform Manifold Approximation and Projection (UMAP). Strains are colour-coded by their taxonomic class to highlight biosynthetic diversity and lineage-specific clustering patterns. The plot reveals distinct groupings, suggesting taxonomically coherent BGC profiles, with some overlap potentially indicating shared ecological niches or horizontal gene transfer.

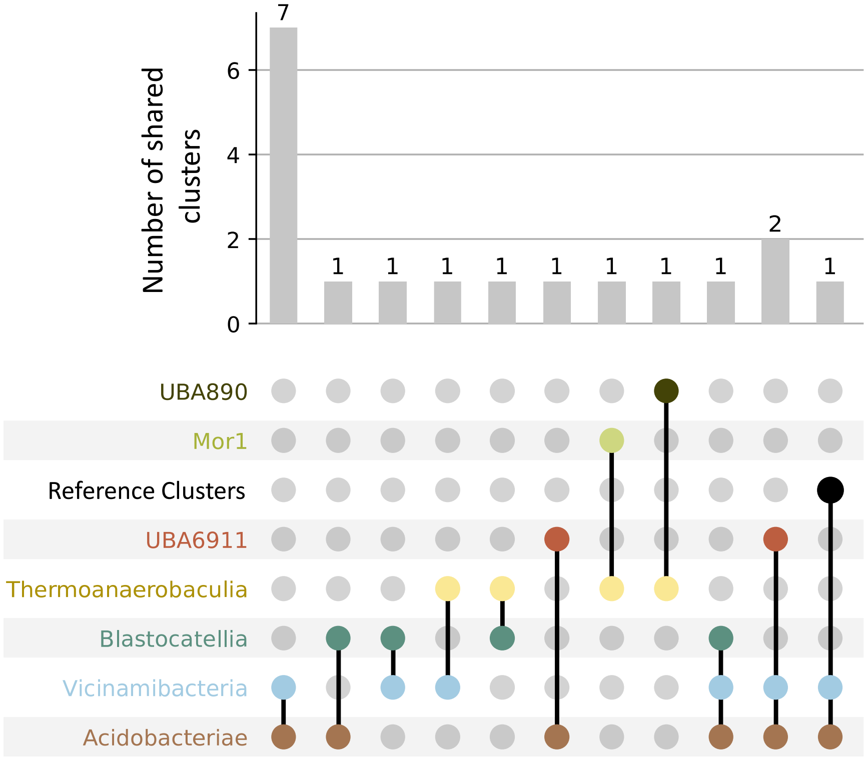


**Figure 10: Gene Cluster Families present in different Acidobacterial Classes.** Bottom: Circles indicate one or more GCFs present in the respective class (left side), lines connect GCFs that are shared between classes. Top: The bar chart indicates the number of GCFs described by the class combinations. The classes *Vicinamibacteria* (blue) and *Acidobacteriae* share the most GCFs (n = 7). Both classes also share two GCFs with the UBA6911 family. Two GCFs are shared between three classes. Both classes are RiPP clusters (data not shown).

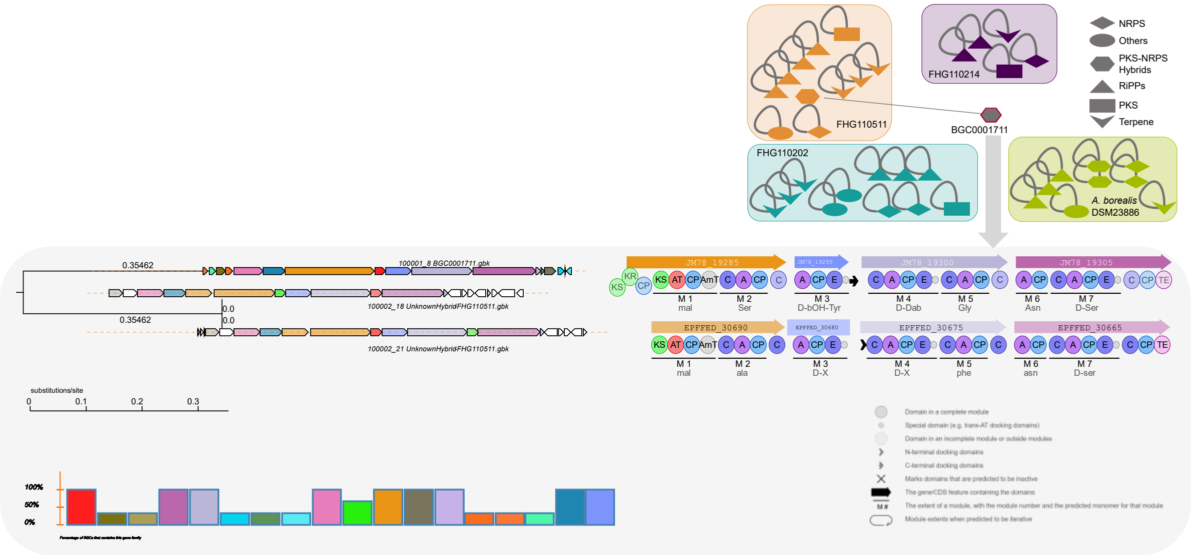


**Figure 11: BiG-SCAPE Similarity Network of antiSMASH-detected BGCs of the FHG Acidobacteriaceae and *A. borealis* DSM 23886 at a cutoff value of 0.5.** MiBIG reference clusters were included to dereplicate similar clusters that had already been described in the literature. The four analysed strains do not share any BGCs and mostly remained as singletons. FHG110511 harbours an NRPS-PKS cluster that shows similarity to the occidiofungin cluster (MiBIG-Reference BGC00001711). Both clusters are PKS-NRPS hybrids, sharing an annotated beta-lactamase gene (red) at a central position within the cluster (pairwise identity: 70.4%). Downstream, both clusters harbour a PKS-NRPS hybrid gene, containing the same number of modules, but not every module matches the a-domain selectivity. Upstream, both clusters contain three multimodular NRPS genes, each harbouring the same number of modules. However, only one gene is predicted to share the amino acid selectivity of the A-domains between the two strains. FHG110511 encodes an additional metallohydrolase within the cluster. At the Upstream border of the cluster, both strains harbour a glycosyl transferase, while downstream we find an additional NRPS-like gene involved in fatty acid synthesis.

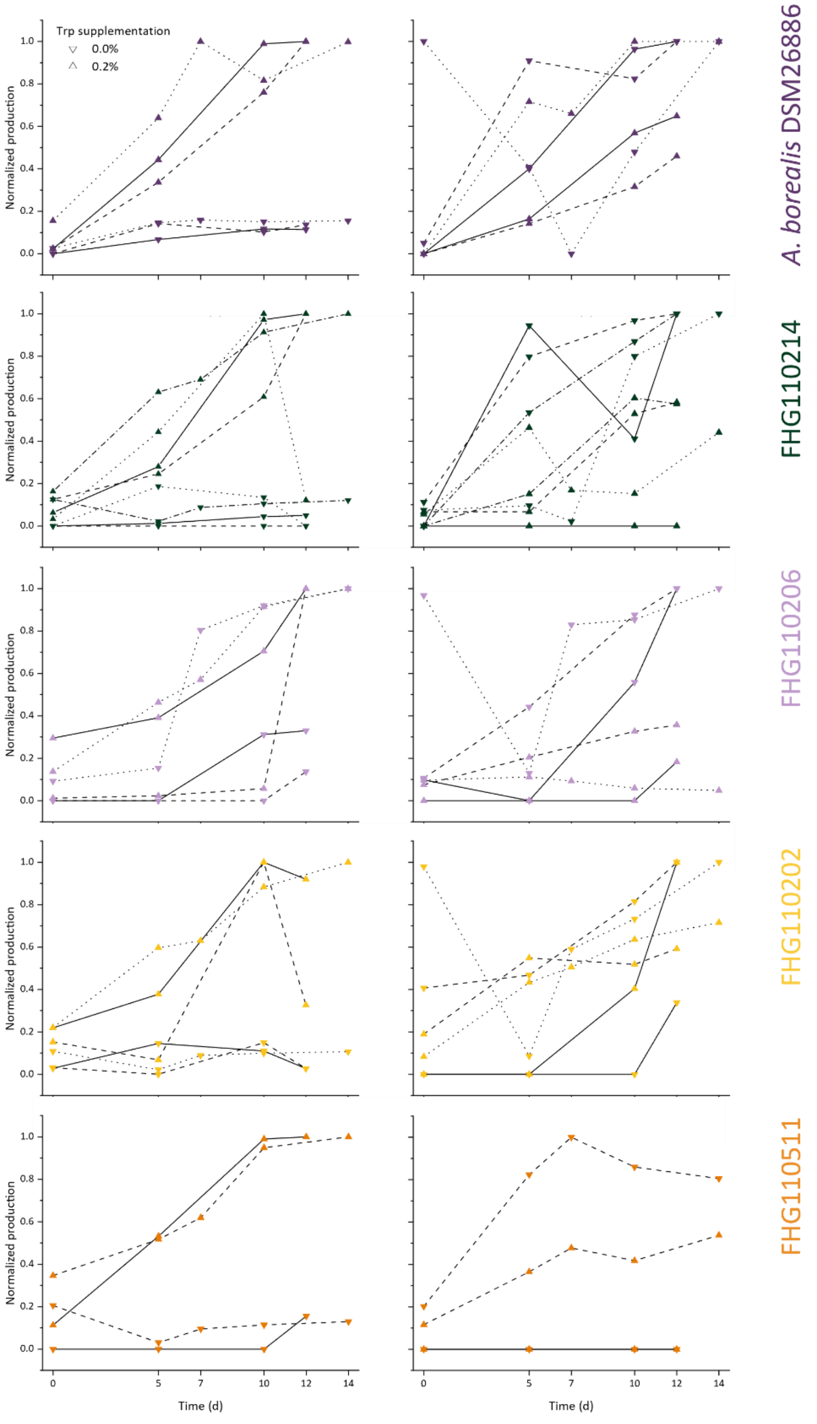


**Figure 12: IAA (left) and iP (right) Production in Investigated Strains Over Time with (triangle) and without Tryptophan (turned triangle).** Available replicates are shown individually to indicate trends. Raw peak areas were normalised within each biological replicate against the highest production of the respective phytohormone. The production of both phytohormones increases over time and typically peaks on the last day of cultivation. While IAA production is consistently increased upon Tryptophan supplementation, the kinetics of iP are less predictable. In general, supplementation with Trp results in a decrease in production, except for strains A. borealis DSM 23886 and FHG110202, where this trend is less pronounced.

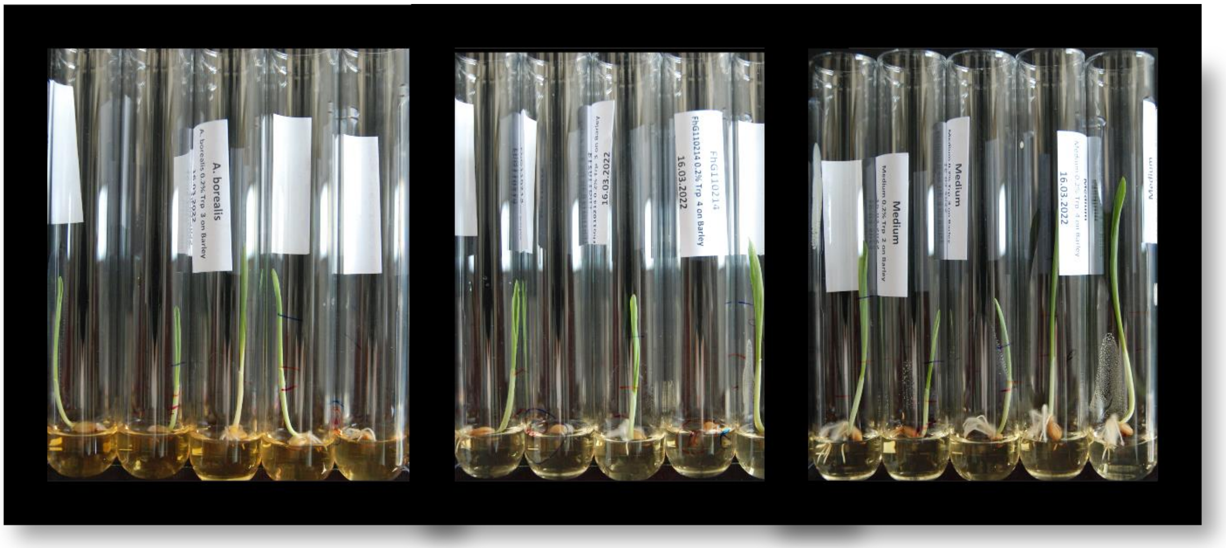


**Figure 13: Experimental setup for plant-growth assays of barley seedlings.** Five replicates of the same condition are shown. Barley seeds are disinfected and grown for 6 days on SH medium that has been pretreated with antibiotics. Extracts from Trp-supplemented cultures were added to the medium as described in the extended Materials and Methods. (A) Extract of *A. borealis* DSM23886 w (B) FHG110214 (C) medium only.

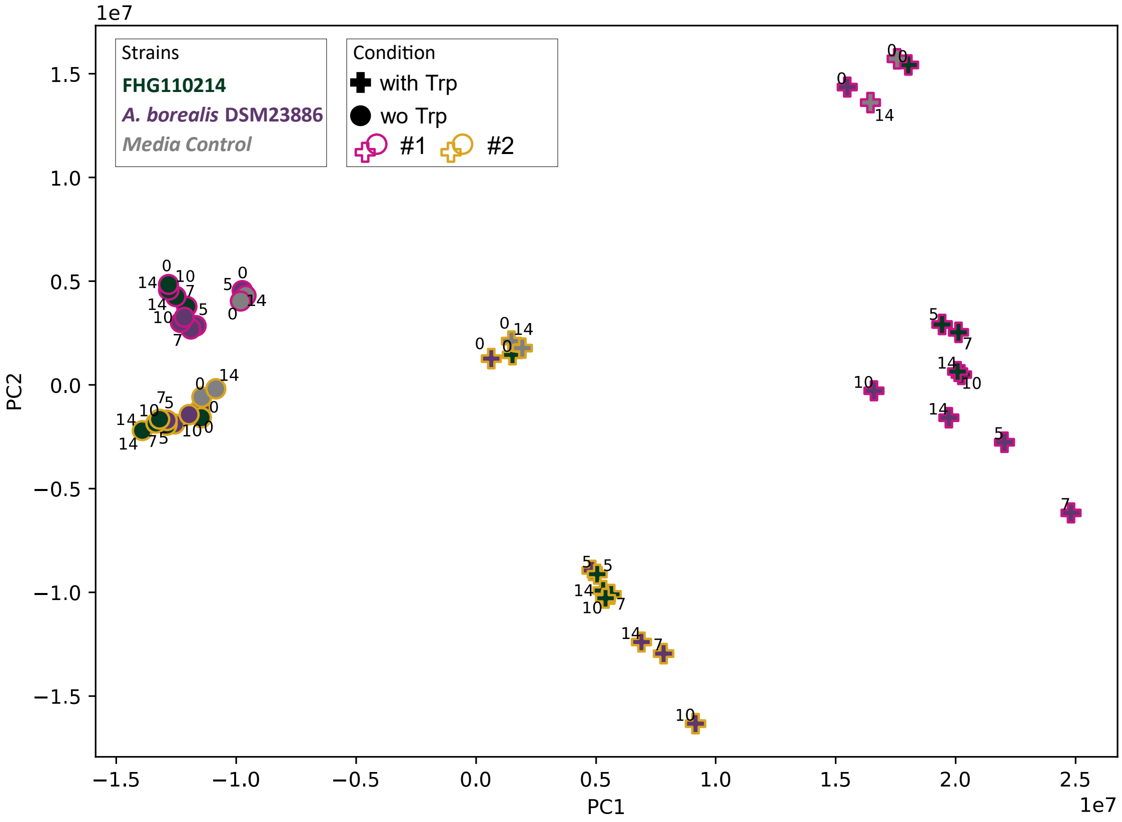


**Figure 14: Principal component analysis of features annotated from crude extracts of A. borealis DSM 23886 and FHG110214 with and without supplementation of Trp.** Biological replicates were normalised to an injection volume of 1 µL before PCA analysis. Border colours indicate biological replicates. Purple colours represent A. borealis DSM 23886, while green colours represent FHG110214. Cross symbols indicate that strains have been supplemented with 0.2% Tryptophan, while circles refer to the uninduced cultures. Media controls are highlighted in grey. All nodes are labelled with their sampling day after cultivation start at day 0. Clusters are grouped by their supplementation status (plus symbols vs. circles) rather than by strain (colours). Biological replicates are shifted but show the same patterns. Notably, extracts generated at the beginning of the cultivation period consistently cluster with the medium controls. The medium controls do not change over the cultivation course. Temporal shifts are most prominent for A. borealis DSM 23886 with Tryptophan induction, than for FHG110214 or A. borealis DSM 23886 without Trp supplementation.

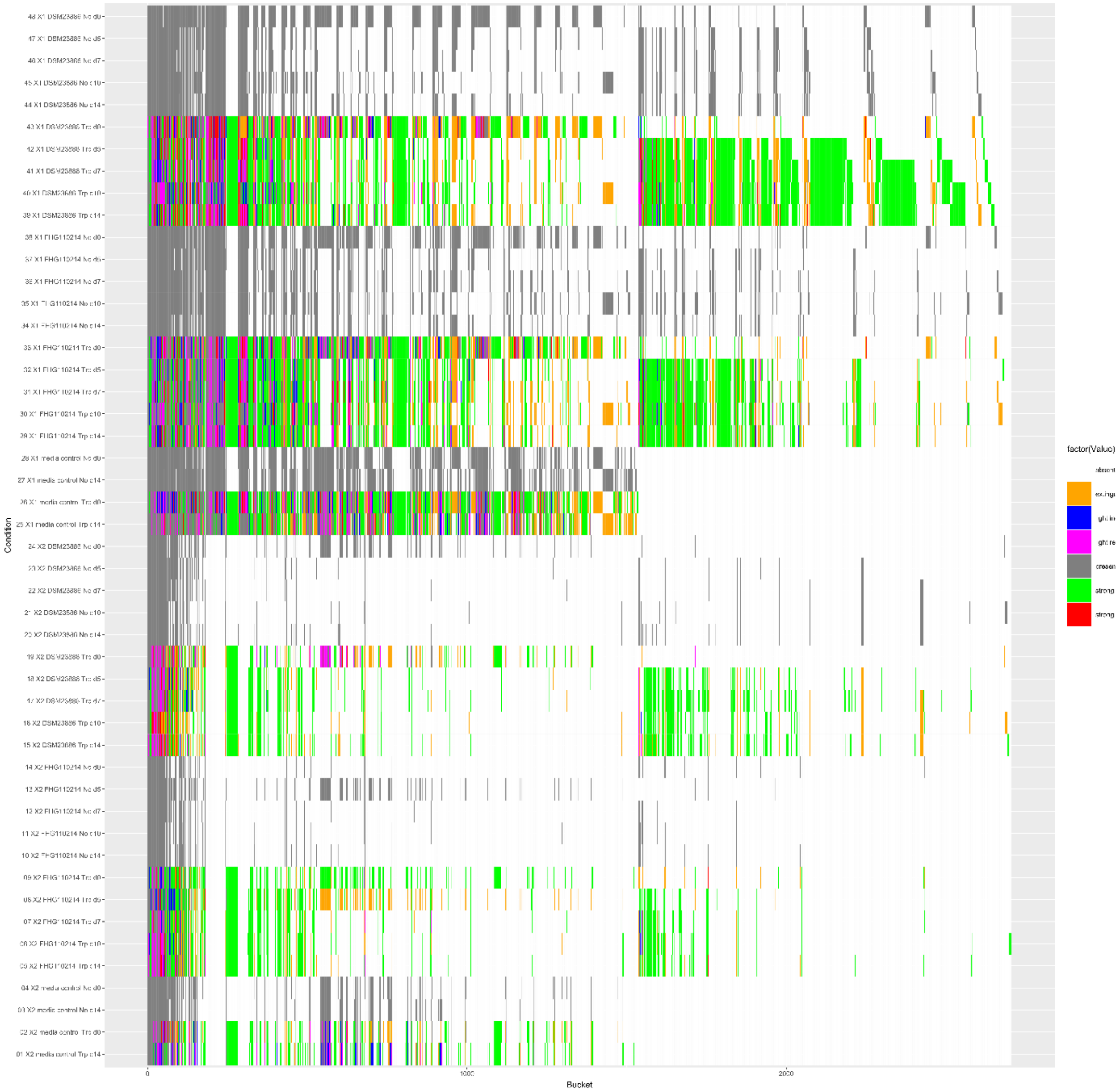


**Figure 15: Chemotype-barcoding upon Trp induction in strain FHG110214 and A. borealis DSM23886**. Untargeted UHPLC-QTOF-HR-MS data were processed by dual-threshold bucketing; only buckets present in the 10,000-threshold set were retained. For the 48 samples, 2759 buckets were calculated. Grey barcode rows represent samples without Trp induction, serving as a reference. Induced samples are classified into response categories compared to the respective uninduced sample: extinguished (orange, indicating absence upon Trp), light increase (blue), light reduction (pink), strong increase (green), and strong reduction (red). Consistent with a global metabolic shift, feature counts increased with Trp (e.g., A. borealis DSM 23886: 165 → 684; FHG110214: 109 → 337).
